# Targeting the 3D genome by anthracyclines for chemotherapeutic effects

**DOI:** 10.1101/2024.10.15.614434

**Authors:** Minkang Tan, Shengnan Sun, Yuchen Liu, Andrea A. Perreault, Douglas H. Phanstiel, Liping Dou, Baoxu Pang

**Author notes:** These authors contributed equally (Minkang Tan, Shengnan Sun, Yuchen Liu). Correspondence (B.P.), (L.D.).

## Abstract

The chromatin is folded into three-dimensional (3D) structures, and aberrant 3D chromatin folding has been implicated in cancer. We performed ATAC-seq and TOP2A ChIP-seq to assess the potential effects of various anthracycline drugs on the chromatin architecture. We found that specific anthracycline variants selectively disrupt chromatin looping anchors by interfering with CTCF binding, suggesting an additional therapeutic mechanism of anthracycline drugs targeting the 3D genome. Hi-C experiments in K562 cells treated with anthracycline drugs revealed widespread disruption of 3D chromatin organization, including altered long-range regulation at the *Myc* locus. Furthermore, AML patients treated with anthracycline drugs exhibited changes in chromatin structures near possible looping anchors, which were associated with distinct clinical outcomes. Together, our findings indicate that anthracycline drugs function as potent and selective epigenomic modulators, with the capacity to further target the 3D genome to exert anticancer effects, highlighting their potential for personalized therapy in tumors with aberrant 3D chromatin architecture.

**Significance:** Chromatin structure plays a crucial role in regulating gene expression and maintaining cellular function. Aberrant chromatin 3D organization is a common feature in cancer. In this study, we investigate how anthracycline chemotherapy affects chromatin architecture using state-of-the-art genomic profiling techniques, including ATAC-seq, TOP2A ChIP-seq, CTCF ChIP-seq, and Hi-C. We show that certain anthracycline variants disrupt chromatin looping by interfering with CTCF binding, thereby altering the spatial genome organization. This disruption leads to changes in the regulation of associated genes, as exemplified by the Myc locus, and is associated with distinct clinical outcomes in AML patients. These findings highlight the potential of anthracycline drugs as therapeutics that target both the epigenome and 3D chromatin architecture, a promising strategy in personalized therapy.

## Introduction

The human genome is packaged into a highly organized structure within the nucleus, forming hierarchical layers of epigenetic regulation that guide biological processes such as cell development and differentiation(1). Emerging techniques such as ChIA-PET, 3C, 4C, and Hi-C, which measure genome-wide contact frequency among genomic loci, have enabled high-resolution studies of nuclear organization(2). Analysis of high-resolution spatial genome data has revealed that chromatin is organized into a three-dimensional (3D) genome architecture, which includes chromosome territories(3), compartments A/B representing transcriptionally active or inactive chromatin(2), topologically associating domains (TADs)(4), architectural stripes(5), and chromatin loops(6). Architectural proteins such as cohesin and CCCTC-binding factor (CTCF) play key roles in mediating 3D genome folding(7). These hierarchical layers of 3D genome structure contribute to the complex regulation of gene transcription within cells.

Growing evidence indicates that cancer cells may have aberrant 3D genome structures, driven by the accumulation of genetic mutations or other mechanisms. These abnormal 3D chromatin interactions could hijack the non-coding regulatory elements, such as silencers(8) and enhancers, to dysregulate gene expression, thereby promoting tumor development and progression(9, 10). These observations highlight the translational and clinical significance of understanding 3D genome architecture in cancer and implicate the potential of targeting the 3D genome for therapeutic intervention. Recently, several studies have shown that small-molecule drugs targeting epigenetic regulators can alter genome organization in tumor cells with diverse effects(9, 11–13). These epigenetic drugs either inhibit transcriptional activators to block enhancer activity(9, 12, 13) or disrupt the liquid-liquid phase separation condensates to interfere with CTCF-independent loop formation(11). However, no currently approved therapeutics have been shown to directly target the 3D genome.

Anthracycline drugs are a class of conventional chemotherapeutic agents that have been widely used in oncology for decades due to their broad-spectrum activity and high efficacy. Therefore, they remain a cornerstone of cancer treatment, particularly for aggressive malignancies such as acute myeloid leukemia (AML). Daunorubicin (Daun) and doxorubicin (Doxo) were among the earliest anthracycline drugs introduced into clinical use. Due to their strong side effects, a variety of anthracycline variants have since been developed, including aclarubicin (Acla), amrubicin (Amr), epirubicin (Epi) and idarubicin (Ida), with distinct therapeutic profiles and tumor-targeting specificities(14). These drugs intercalate into the DNA and most of them induce DNA double-strand breaks (DSBs) by interfering with the catalytic cycle of topoisomerase IIα (TOP2A)(15). More recently, anthracycline drugs have been shown to exert anticancer effects via a novel mechanism—histone eviction from chromatin—inducing tumor cell death independently of DNA damage(16). Notably, the anthracycline variant Acla, used clinically to treat AML has been demonstrated to evict histones without causing DSBs(16, 17) and to lead to RNA Pol II degradation(18). Moreover, different anthracycline variants may selectively target distinct genomic regions, contributing to their varied therapeutic effects(16, 19).

In this study, we show that anthracycline drugs, exemplified by Acla and Daun, redistributed their canonical target protein TOP2A around chromatin looping anchors, as shown by ChIP-seq of endogenously tagged TOP2A in human erythroleukemia K562 cells. ATAC-seq analysis revealed reduced nucleosome compaction and diminished CTCF footprint signals at CTCF-bound looping anchor regions, which are typically associated with the formation of insulating chromatin domain boundaries(20). This reduction of CTCF binding was further validated by western blotting and CTCF ChIP-seq, indicating that these drugs may impact 3D chromatin architecture. Indeed, in situ Hi-C experiments revealed widespread spatial genome alterations following Acla and Daun treatment in K562 cells, including weakened TAD boundary insulation and disruption of chromatin loops. Integrated analyses combining in situ Hi-C, ATAC-seq, and ChIP-seq suggest that reduced CTCF occupancy may be a primary driver of loop decoupling. To assess whether anthracycline treatment also induces 3D chromatin changes in patients, we performed ATAC-seq on tumor cells from AML patients treated with Daun, which revealed that CTCF-bound regulatory elements were similarly affected. Furthermore, we observed that the potential 3D chromatin changes in AML patient tumor cells correlated with patient clinical outcomes following anthracycline-based chemotherapy. These findings indicate that anthracycline drugs may also exert therapeutic effects by targeting 3D chromatin structure, a property that could potentially be harnessed to treat other cancers characterized by aberrant chromatin organization.

## Results

### Anthracycline drugs redistribute TOP2A at chromatin looping anchors

Most anthracycline drugs exert their cytotoxic effects by inhibiting TOP2A activity(15). In addition to its role in DNA replication and repair, TOP2A is also involved in transcriptional regulation and genome organization(21). To examine the impact of anthracycline drugs on the interaction between TOP2A and chromatin, we performed genome-wide mapping of TOP2A binding using chromatin immunoprecipitation followed by sequencing (ChIP-seq) in K562 cells with endogenously FLAG-tagged TOP2A (see **Methods**). Previous studies have shown that promoter-associated TOP2A helps resolve transcription-induced DNA supercoiling(21, 22). Consistent with this, we observed TOP2A accumulation at promoter regions, with its distribution altered following treatment with aclarubicin (Acla) or daunorubicin (Daun) (**Supplementary Fig. 1a and b**). While TOP2A shares partial functional redundancy with TOP2B(23), which was shown to interact with CTCF and cohesin at TAD boundaries to relieve topological stress during loop extrusion(21, 24, 25), our data showed that Acla and Daun treatment led to a marked reduction of TOP2A binding at TAD boundaries co-occupied by CTCF and cohesin. This effect was not observed with amrubicin (Amr), an anthracycline variant that induces DNA damage but does not cause histone eviction (**Fig. 1a**). Further analysis of all chromatin looping anchors bound by CTCF and cohesin revealed a clear reduction in TOP2A ChIP-seq signal upon Acla and Daun treatment. Daun treatment led to a marked depletion of TOP2A ChIP-seq signals primarily at CTCF/cohesin looping anchors, whereas Acla significantly reduced TOP2A ChIP signals within a broader ±2 Kbp window around these anchors. These differences likely stem from the distinct chemical structures and epigenomic targeting properties of the two drugs (**Fig. 1b**). To quantitatively assess TOP2A redistribution, we performed differential binding analysis and found that Acla and Daun caused widespread perturbations in TOP2A occupancy across the genome, whereas Amr had minimal impact compared to untreated controls (**Supplementary Fig. 1c**). Permutation-based association testing revealed that lost or stably maintained TOP2A peaks following Acla and Daun treatment were significantly associated with CTCF-binding sites, while newly gained TOP2A peaks showed no such association (**Fig. 1c**). Motif enrichment analysis of lost TOP2A peaks further identified significant enrichment for CTCF, CTCFL, and GATA2 motifs (**Fig. 1d**), suggesting that Acla and Daun disrupt TOP2A binding at looping anchors and TAD boundaries marked by these factors by CTCF. Interestingly, this pattern was also observed with other clinically used anthracycline drugs known to induce histone eviction(17) (**Supplementary Fig. 1d**). Together, these results suggest that many anthracycline drugs may promote the redistribution of TOP2A away from CTCF-associated TAD boundaries and chromatin looping anchors.

**Figure 1.**
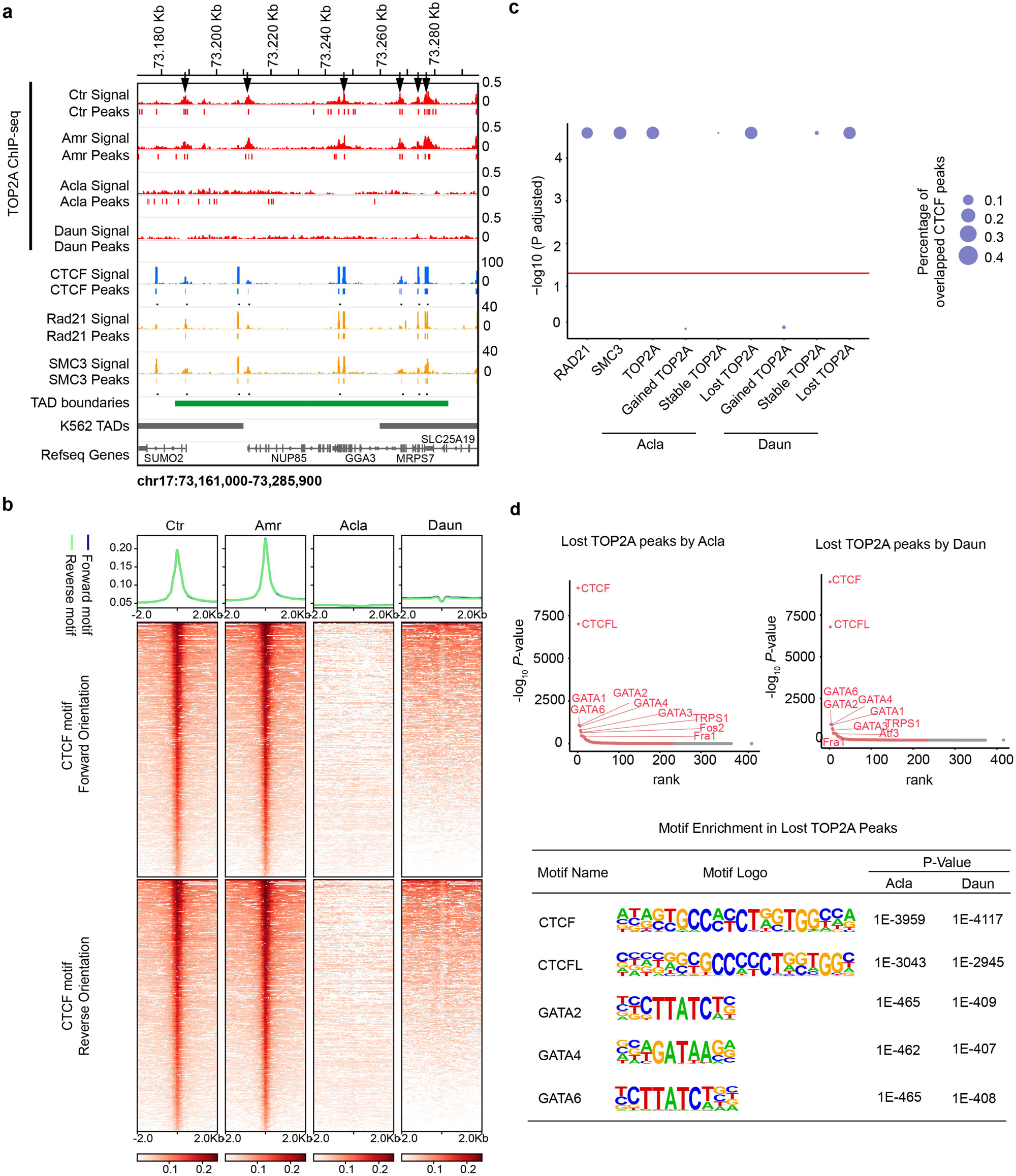
Anthracyclines redistribute TOP2A at chromatin looping anchors. **(a)** Snapshot of TOP2A ChIP-seq data from untreated (Ctr), Amr-, Acla- and Daun-treated K562 cells. K562 cells were treated with 10□µM of the respective drugs for 2 hours. ChIP-seq against CTCF, cohesin subunits RAD21 and SMC3, TADs, TAD boundaries, and gene annotation in K562 cells are shown. **(b)** TOP2A ChIP-seq profiles at chromatin loop anchors in untreated (Ctr), Amr-, Acla-, and Daun-treated K562 cells. Chromatin looping anchors were defined by co-binding of CTCF and RAD21 (cohesin) based on ChIP-seq data. CTCF motifs within anchors were identified using FIMO (*P*-value < 10^-5^) and categorized as forward or reverse orientation based on detected DNA strand. Normalized TOP2A ChIP-seq signals are shown ±2 kb from the CTCF motif center. **(c)** Association of TOP2A peaks in untreated (Ctr) K562 cells and differential TOP2A peaks from Acla- and Daun-treated cells with CTCF ChIP-seq peaks. RAD21 and SMC3 (cohesin subunits) serve as positive controls for CTCF association. The y-axis shows enrichment significance of TOP2A within CTCF peaks, calculated by permutation testing (*n*=50,000) with Benjamini-Hochberg correction. The red horizontal line marks the significance threshold (*P*-value=0.05). Circle size reflects the proportion of CTCF peaks overlapped by each TOP2A peak set (scale: 0-1). **(d)** TF motif enrichment in lost TOP2A peaks from Acla- and Daun-treated K562 cells calculated by HOMER. The top panel shows the ranked list of enriched TF motifs. The bottom panel displays the sequence logos and *P*-values of the top five enriched motifs.

### Anthracycline treatments affect CTCF footprinting on chromatin

Recent studies have shown that anthracycline drugs selectively evict histones from defined chromatin structures(16, 17, 19). We hypothesized that anthracycline-induced depletion of TOP2A from chromatin looping anchors marked by CTCF may coincide with direct disruption of CTCF binding. To investigate this, we used ATAC-seq to profile chromatin accessibility and infer genome-wide changes in transcription factor (TF) binding following anthracycline treatment. K562 cells were treated with Acla, Amr or Daun for 2 hours before ATAC-seq. Cells treated with histone-evicting anthracycline drugs (Acla and Daun) showed a distinct fragment size distribution compared to Amr-treated (inducing DNA damage only) and untreated cells (**Fig. 2a**), consistent with altered nucleosome periodicity due to histone eviction(26). In addition, Tn5 insertion patterns were significantly altered in a state-specific manner across seven previously defined functional chromatin states(27) (**Fig. 2b**). In particular, CTCF-bound regions, which are typically enriched for shorter fragments and depleted of longer ones due to CTCF-mediated nucleosome positioning(28, 29), exhibited altered nucleosome organization following Acla and Daun treatment (**Fig. 2b**). ChromHMM-based analysis of 15 chromatin states corroborated these findings, showing notable changes in insulator elements(30) (**Supplementary Fig. 2a-d**). We then performed nucleosome occupancy profiling at CTCF-bound looping anchor regions using NucleoATAC(31). This analysis revealed a substantial decrease in nucleosome occupancy at these sites following Acla and Daun treatment (**Fig. 2c**), which may reflect weakened chromatin insulation at domain boundaries(20). We further analyzed nucleosome positioning around transcription start sites (TSSs) and observed disruption of the nucleosome-free regions flanked by well-positioned nucleosomes, supporting a global chromatin remodeling effect of anthracycline drugs(32) (**Supplementary Fig. 2e**). Finally, to unbiasedly assess transcription factor binding changes, we performed genome-wide CTCF footprinting analysis using HINT-ATAC(33). Aggregated footprints at CTCF motifs showed that Acla and Daun treatments diminished CTCF occupancy, likely through disruption of local chromatin structure(29) (**Fig. 2d**). Together, these data suggest that Acla and Daun altered the chromatin landscape around looping anchors and weakened CTCF binding.

**Figure 2.**
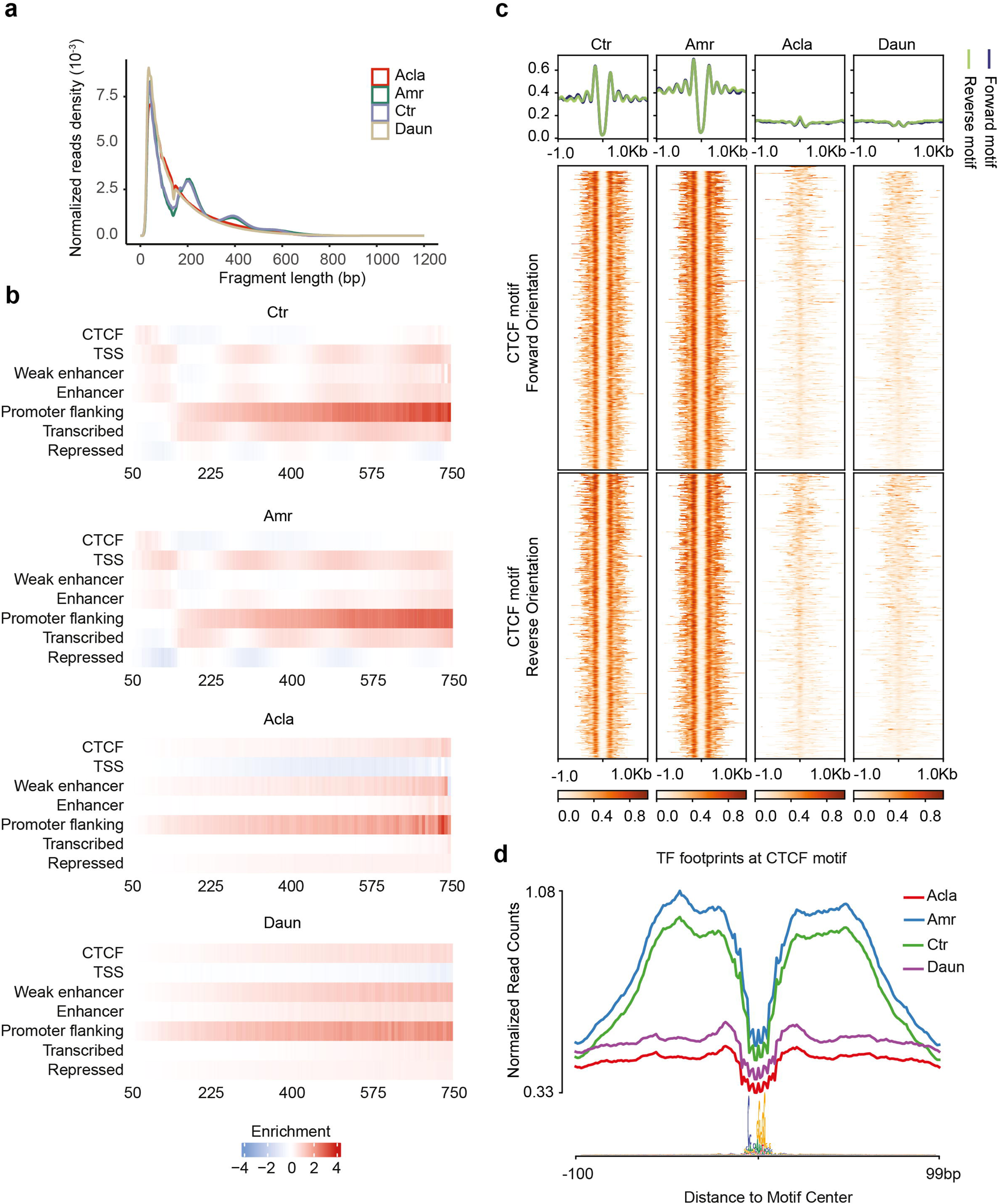
Anthracyclines affect CTCF foot-printing on chromatin. **(a)** ATAC-seq fragments size distribution from untreated (Ctr), Amr-, Acla- and Daun-treated K562 cells. K562 cells were treated with10□µM of respective drugs for 2hours. **(b)** Normalized ATAC-seq fragment enrichment in seven chromatin state classes from untreated (Ctr), Amr-, Acla-, and Daun-treated K562 cells. Chromatin states were defined in K562 cells as follows: CTCF (CTCF-bound elements), TSS (promoter regions with transcription start sites), Weak enhancer (putative weak enhancers/open chromatin), Enhancer (predicted enhancers), Promoter flanking (promoter-adjacent regions), Transcribed (actively transcribed regions), and Repressed (low-activity or repressed regions). Tn5 insertion fragments (50–750 bp) were divided into 100 bins, and insertion percentages per bin were normalized to the maximum within each state. Enrichment was calculated by dividing the normalized percent of insertions for each state by the genome-wide normalized percent of insertions. Scale: –4 to +4. **(c)** Nucleosome occupancy around chromatin loop anchors in untreated (Ctr), Amr-, Acla-, and Daun-treated K562 cells. Loop anchors were defined as co-binding sites of CTCF and RAD21 (cohesin) from ChIP-seq data. CTCF motifs were identified using FIMO (*P*-value<10^-5^) and classified by strand orientation. Nucleosome occupancy is shown ±1 kb from the CTCF motif center. **(d)** TF footprinting around CTCF motifs (n=16,814) based on ATAC-seq data from untreated (Ctr), Amr-, Acla-, and Daun-treated K562 cells. Normalized Tn5 cleavage frequency is shown ±100 bp from the CTCF motif center.

### Anthracycline drugs evict CTCF from chromatin

To directly assess whether anthracycline drugs affect CTCF binding, K562 cells were treated with 10μM Daun for 2 hours, followed by cellular fractionation to separate the non-chromatin and chromatin-bound proteins. Immunoblotting revealed a reduction of CTCF in the chromatin-bound fraction, accompanied by an increase in the non-chromatin fraction, compared to untreated cells, indicating that Daun may partially evict endogenous CTCF from chromatin (**Fig. 3a and b**). Total CTCF protein levels did not change, suggesting that the shift in CTCF localization was not due to overall protein loss. To further examine CTCF disruption on a genome-wide scale, we performed CTCF ChIP-seq following treatment with Acla or Daun. The results showed widespread loss of CTCF binding across the genome (**Fig. 3c**). Interestingly, we also detected some new CTCF-binding sites after drug treatment (**Supplementary Fig. 3a and b**), suggesting that anthracycline drugs may promote chromatin looping reorganization by both disrupting and relocating CTCF binding. Given the central role of CTCF in organizing chromatin looping and TADs(34, 35), we next examined its occupancy at these genomic structures. CTCF ChIP-seq signal intensity decreased globally at looping anchors and TAD boundaries following Acla and Daun treatment (**Fig. 3d, upper panels**). Notably, CTCF binding near TSSs, which often defines enhancer- or silencer-promoter interactions(36, 37), was also reduced upon Acla and Daun treatment (**Fig. 3d, lower panels**). These observations suggest that anthracycline drugs may disrupt cis-regulatory landscapes by interfering with CTCF-mediated interactions. As TOP2A was previously shown to be redistributed from chromatin looping anchor regions and TAD boundaries (**Fig. 1**), we further investigated regions where CTCF and TOP2A colocalize(24, 38). Integrative analysis of TOP2A and CTCF ChIP-seq data revealed co-depletion of both proteins from these shared sites following Acla and Daun treatment (**Fig. 3e**). Together, these findings demonstrate that anthracycline treatment may disrupt CTCF binding on chromatin, potentially contributing to large-scale reorganization of 3D genome architecture.

**Figure 3.**
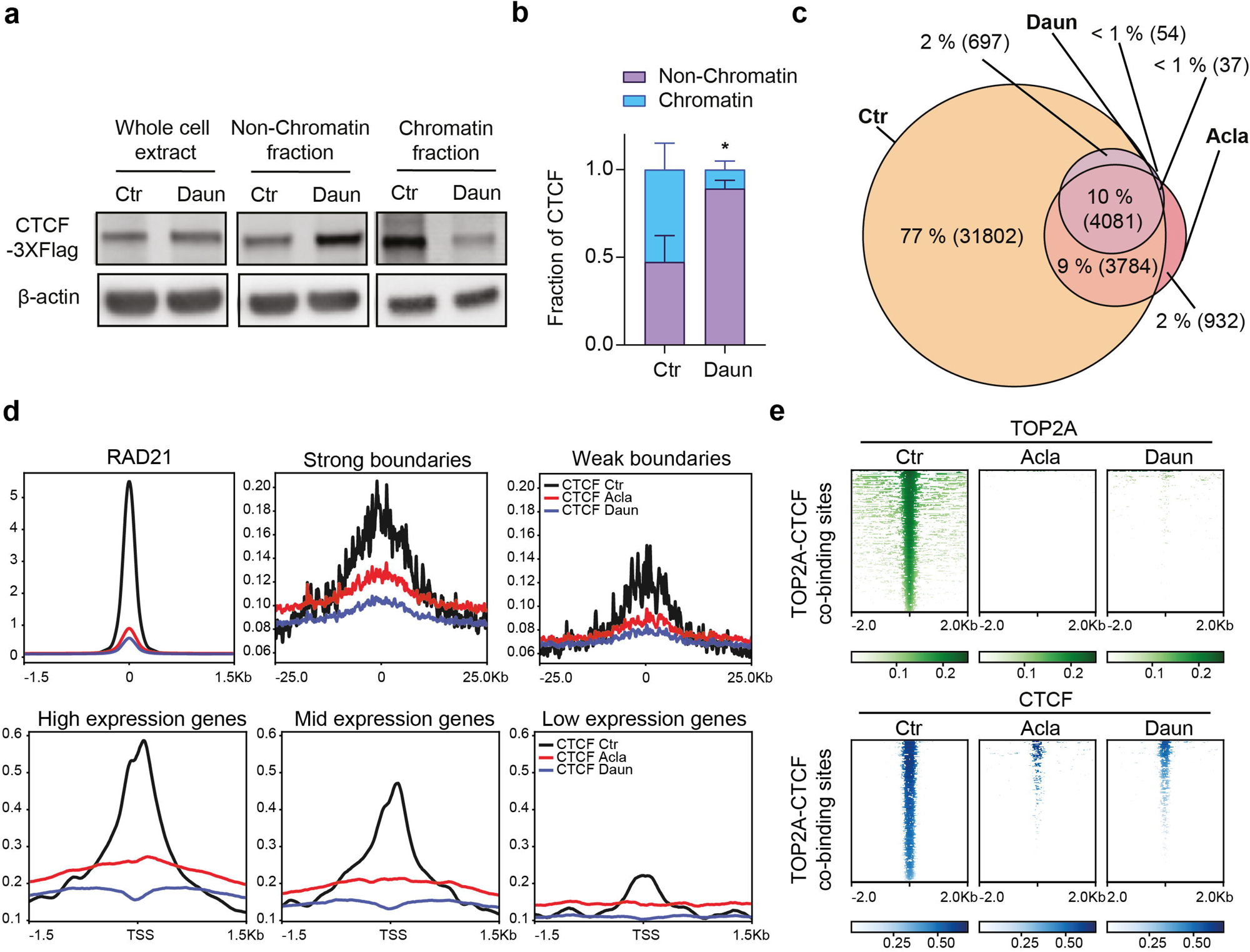
Anthracyclines evict CTCF from chromatin. **(a)** CTCF protein distribution in the whole cell extract, non-chromatin and chromatin fraction in untreated (Ctr) and Daun-treated K562 cells by western blot. The K562 Cells with endogenously 3×FLAG-tagged CTCF were treated with 10□µM Daun for 2 hours; untreated cells served as the control. β-actin was used as the loading control. **(b)** Quantification of CTCF levels in each cellular fraction from Western blot analyses across three biological replicates. \**P*=0.0161, calculated by two-way ANOVA. **(c)** Venn diagram of CTCF peaks identified from ChIP-seq in untreated (Ctr), Acla-, and Daun-treated K562 cells. Cells were treated with 10□µM Acla or Daun for 2 hours; untreated cells served as the control. Two biological replicates were analyzed per condition. **(d)** Density plots of CTCF ChIP-seq signals at cohesin RAD21 binding sites, TAD boundaries, and transcription start sites (TSSs) in untreated (Ctr), Acla-, and Daun-treated K562 cells. TAD boundaries were identified from published K562 in situ Hi-C data using the diamond insulation method at 10 kb resolution. Strong boundaries were defined as local minima with log₂(insulation score) > 0.5; weak boundaries, as those between 0.2 and 0.5. TSSs were grouped into high-, medium-, and low-expression categories based on K562 RNA-seq data. **(e)** TOP2A and CTCF redistribution at TOP2A-CTCF co-binding sites upon Acla and Daun treatment. Co-binding regions were defined from CTCF and TOP2A ChIP-seq data in untreated K562 cells. Normalized TOP2A (top) and CTCF (bottom) ChIP-seq signals are shown ±2 kb around the interaction peak center.

### Anthracycline drugs impact 3D genome organization

To test the hypothesis that anthracycline treatment impacts 3D genome architecture, K562 cells treated with Acla or Daun were subjected to *in situ* Hi-C to assess chromatin folding changes (**Fig. 4a**). A substantial increase in the trans-to-cis contact ratio was observed after drug treatment (**Supplementary Fig. 4a**), based on high quality Hi-C datasets (**Supplementary Table 2**). Analysis of intrachromosomal interactions revealed a steep decay in short-range contacts (<10□kb) and an increase in long-range contacts (>1□Mb) following anthracycline exposure (**Fig. 4b**). Moreover, saddle plot analysis showed weakened compartmentalization (**Fig. 4c**), and TAD boundary strength was reduced as measured by the diamond insulation method(39) (**Fig. 4d**). To detect differentially insulated TAD boundaries induced by drug treatment, we first calculated insulation scores at 10-kb resolution and identified boundaries based on differences from untreated controls(40). The analysis revealed that most TAD boundaries were lost following anthracycline treatment (73.97% with Acla and 81.52% with Daun) (**Fig. 4e**). These losses were consistent with CTCF depletion at chromatin boundaries (**Fig. 3**), and mirrored the effects previously reported for RNAi-mediated CTCF knockdown, which led to a 63.83%reduction in boundaries(41)(**Fig. 4e**). To assess the impact on the remaining TAD boundaries (including the weakened, stable, and enhanced boundaries identified in **Fig. 4f**), we analyzed correlations in boundary strength scores (**Fig. 4f, Supplementary Fig. 4b**) and insulation profiles (**Supplementary Fig. 4c and d**). The results confirmed reduced insulation strength at residual boundaries following treatment. We further investigated whether the TAD boundary disruptions observed with Acla or Daun could be attributed to CTCF loss (**Supplementary Fig. 4e and f**). Comparison of Daun-treated cells with CTCF siRNA knockdown cells revealed that 55.1% of the disrupted boundaries were shared. Daun alone disrupted an additional 26.4% of boundaries, while CTCF knockdown uniquely affected 8.7%. The remaining 9.7% of boundaries were unaffected under either condition (**Supplementary Fig. 4f**). Together, these findings indicate that anthracycline drugs may disrupt 3D genome architecture by evicting CTCF from its native chromatin context.

**Figure 4.**
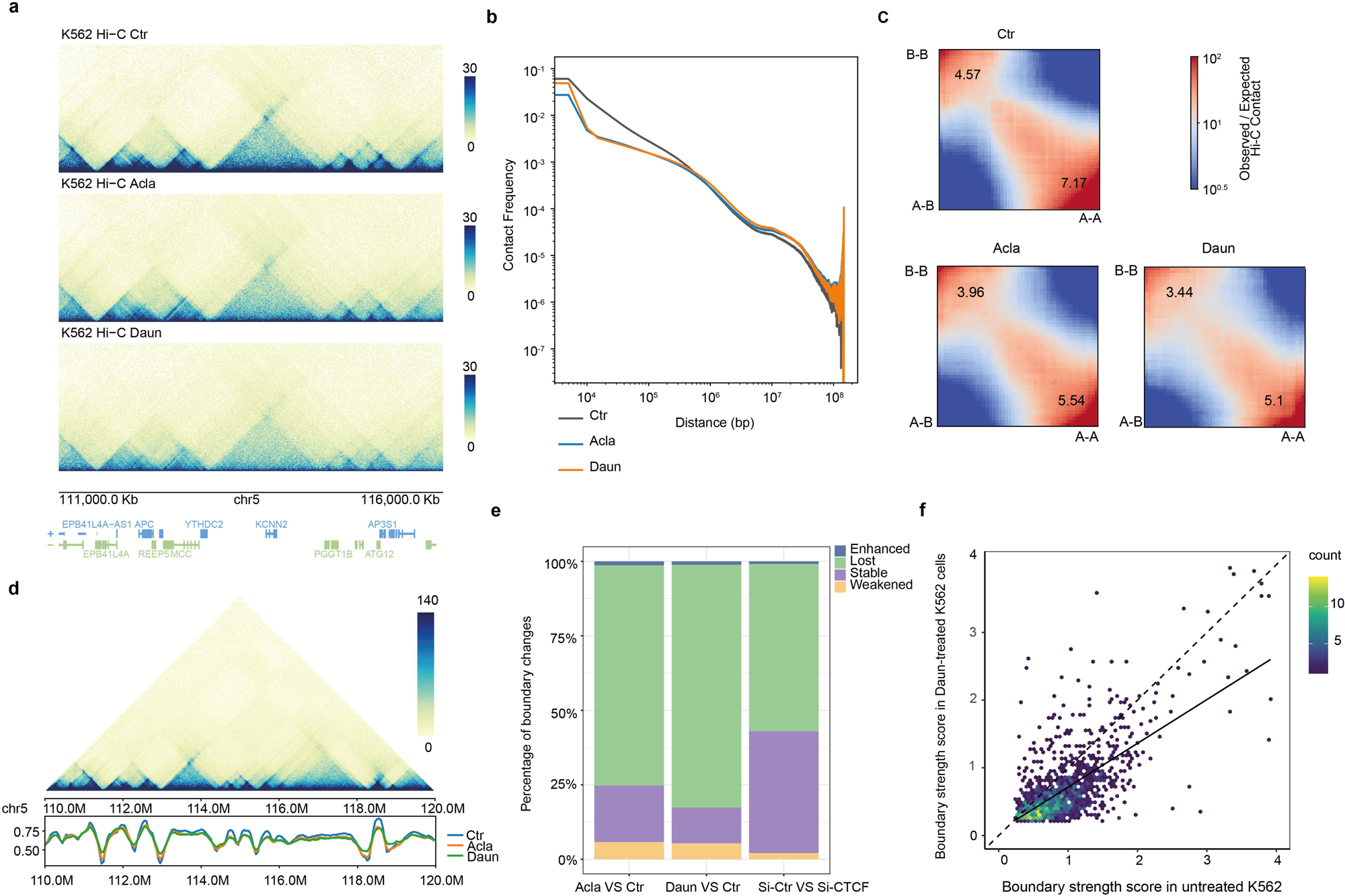
Anthracyclines impact 3D genome organization. **(a)** Snapshot of Hi-C maps. From top to bottom, Hi-C matrices of untreated (Ctr), Acla- and Daun-treated K562 cells. Hi-C data analysis was performed on K562 cells treated with Acla or Daun at 10□µM for 2 hours. Untreated cells served as controls (Ctr). **(b)** Decay curves of chromatin interaction frequency with distance in untreated (Ctr), Acla- and Daun-treated K562 cells. The *y*-axis represents the logarithm scale of contact frequency from respective Hi-C data and *x*-axis represents logarithm scale of the genomic distance between looping anchors (bp). **(c)** Saddle plots showing A/B compartment separation strength in untreated (Ctr), Acla-, and Daun-treated K562 cells (left to right). Average contact frequencies between 100 kb bins were normalized by expected contact frequency based on genomic distance. Bins were ranked by first eigenvector values and divided into 100 quantiles. Red areas in the upper left and lower right indicate enriched B-B and A-A interactions, respectively, while blue areas in the upper right and lower left show depleted B-A and A-B interactions. Numbers in the corners represent the relative strengths of A-A versus A-B (lower right) and B-B versus B-A (upper left) interactions. **(d)** Snapshot of insulation score tracks derived from KR-normalized Hi-C matrices (10 kb bins, 100 kb window) using the diamond-insulation algorithm. Upper panel: Hi-C contact map at chr5:111,000,000–116,000,000. Lower panel: insulation profiles for untreated (Ctr), Acla-, and Daun-treated K562 cells, color-coded accordingly. Insulation scores reflect contact frequencies between upstream and downstream regions; local minima of insulation scores define boundary bins. **(e)** Changes in insulation strength at TAD boundaries in Acla-, Daun-, and si-CTCF-treated K562 cells. Four boundary classes are shown: lost (absent in drug-treated Hi-C), weakened (control-drug boundary score >= 0.4 and control-drug insulation score <= -0.1), stable, and enhanced (control-drug boundary score <= -0.4 and control-drug insulation score >= 0.1). Genome-wide insulation and boundary scores were calculated from KR-normalized Hi-C matrices (10 kb resolution, 100 kb window) using the diamond-insulation algorithm. Published Hi-C from si-CTCF-treated K562 cells (48-hour treatment) were included for comparison. **(f)** Scatterplot of boundary strength scores for residual TAD boundaries from untreated (x-axis) and Daun-treated (y-axis) Hi-C matrices. The solid line shows the regression fit; the dashed diagonal line indicates equality. The color indicates the spatial density of nearby points.

### Anthracycline drugs affect CTCF-mediated *cis*-regulatory loops

To investigate whether chromatin looping structures within TADs were affected by anthracycline drugs, we further analyzed Hi-C data from Acla- and Daun-treated K562 cells using the random-forest-based classifier Peakachu(42), which accounts for sample-specific sequencing depth variation. A Gaussian mixture model was applied to identify differential loops upon drug treatment, and the results were visualized by aggregate peak analysis (APA)(10)(**Fig. 5a, Supplementary Fig. 5a**). This analysis revealed substantial disruption of existing chromatin loops following drug treatment (24,460 and 28,025 loops were lost in Acla- and Daun-treated cells, respectively, FDR < 0.05), while some new loops were also gained (6,328 and 4,800 loops were gained in Acla- and Daun-treated cells, respectively, FDR < 0.05). Within the stable chromatin loops, the local contact background was reduced in drug-treated samples compared to controls, likely reflecting drug-induced disruption of nearby chromatin interactions of the stable loops. Integrative analysis of ATAC-seq and ChIP-seq data showed that CTCF was one of the most significantly associated factors at these lost loops (**Fig. 5b, Supplementary Fig. 5b**). Specifically, loss of CTCF ChIP-seq peaks coincided with loss of ATAC-seq peaks containing CTCF motifs during drug treatment, suggesting that loop disruption was driven by CTCF eviction from chromatin (**Fig. 3**), consistent with its known role in loop formation and stabilization(43). Notably, more stable CTCF peaks, as defined by CTCF ChIP-seq, persisted at lost looping anchors following Acla treatment (**Supplementary Fig. 5b**). Given that chromatin loops are often stabilized by clusters of redundant CTCF binding to ensure robust long-range interactions, it is likely that Acla selectively evicts a subset of CTCF from these clusters. This partial loss could weaken looping robustness without completely abolishing CTCF binding at these regions. Interestingly, the distance between anchors of newly formed loops in Acla- and Daun-treated cells was significantly longer than lost or unchanged stable loops (**Supplementary Fig. 5c and d**). To assess the functional consequences of disrupted loops by these drugs, we performed KEGG pathway analysis on genes whose TSSs were associated with lost loops following drug treatments. The results implicated pathways related to the cell cycle, Wnt signaling, and Hippo signaling as targets of anthracycline-mediated disruption (**Supplementary Fig. 5e and f**). Among the drug-disrupted loops, we also observed loss of well-established CTCF-mediated promoter–enhancer and enhancer–enhancer interactions that regulate *Myc* overexpression in K562 cells(44, 45) (**Fig. 5c**). Differential Hi-C loops and virtual 4C analysis anchored at the *Myc* promoter revealed disrupted interactions with *Myc* enhancers following treatment (**Fig. 5d**), consistent with decreased CTCF binding at these sites. Moreover, enhancer–enhancer interactions within the *Myc* regulatory landscape were also lost, potentially impairing the cooperative enhancer network that ensures robust *Myc* expression(45) (**Fig. 5c**). These findings were further supported by additional 4C-seq experiments, which showed disrupted distal interactions at the *Myc* locus and other regulatory regions, such as the human β-globin locus, following Daun treatment (**Supplementary Fig. 5g-i**). To broaden this observation, we evaluated the effects of Acla and Daun on five major categories of cis-regulatory looping interactions (CTCF–CTCF, promoter–promoter, promoter– enhancer, enhancer–enhancer, and putative silencer–promoter loops) via APA analysis(46). The results demonstrated widespread disruption of these interactions genome-wide (**Fig. 5e, Supplementary Fig. 5j**). Together, these data suggest that anthracycline drugs have promising therapeutic potential by targeting aberrant cis-regulatory chromatin loops, particularly those mediated by CTCF, which drive oncogene activation or repress tumor suppressor genes(47).

**Figure 5.**
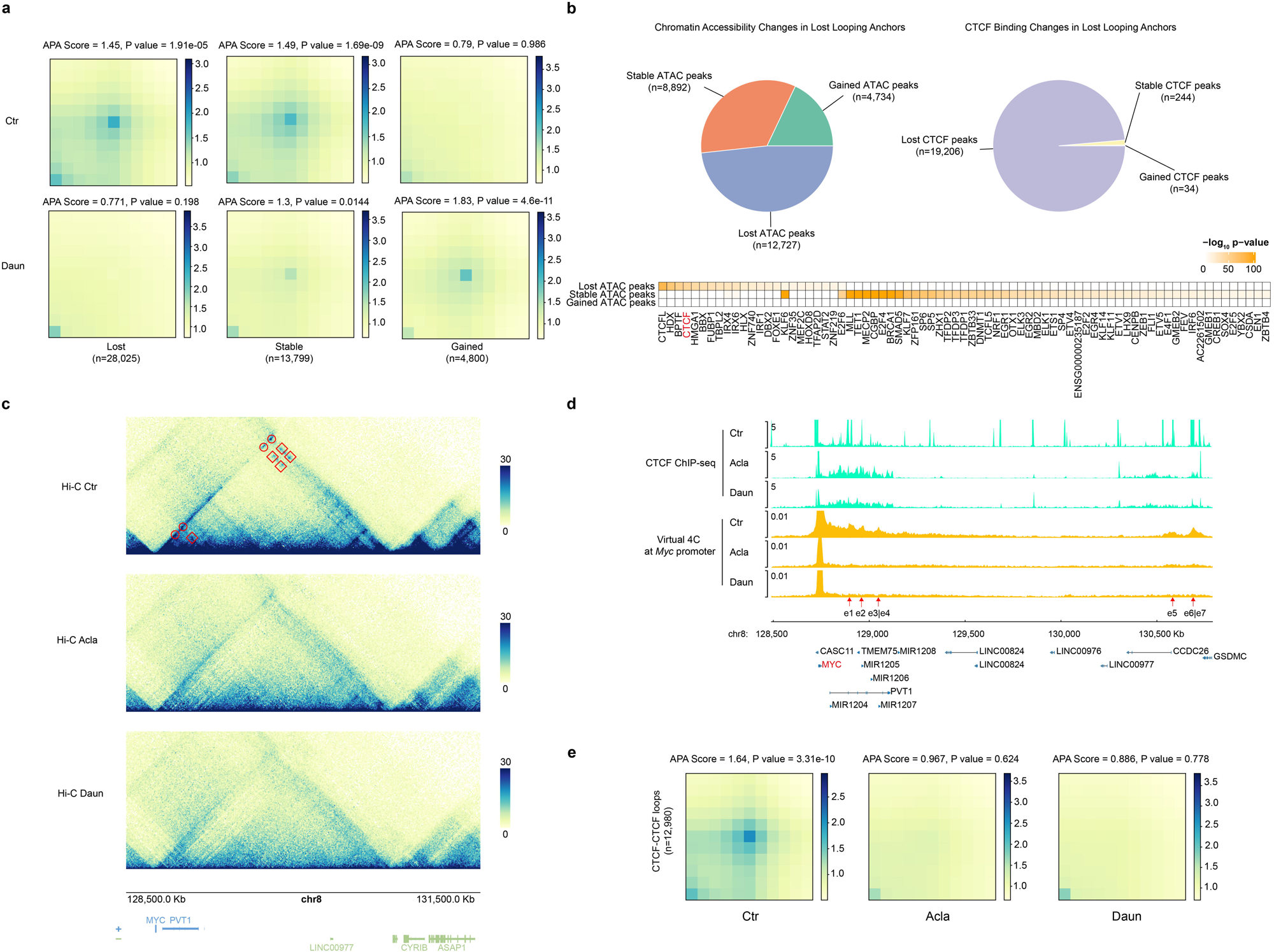
Anthracyclines affect CTCF-mediated *cis*-regulatory loops. **(a)** Aggregate peak analysis of rewired chromatin loops in Daun-treated K562 cells (gained, lost and stable) within ± 50 Kbp window. Differential loops were determined by diffPeakachu when the fold-change of probability score deviates from the Gaussian mixture model with a P-value□<□0.05. **(b)** Epigenomic signatures of lost loops in Daun-treated K562 cells. Upper panel: pie charts of Daun-induced differential (gained, lost and stable) ATAC-seq peaks (left) and CTCF ChIP-seq peaks (right) within the lost looping anchors. Lower panel: motif enrichment in Daun-induced differential ATAC-seq peaks (from top to bottom: lost, stable and gained). *P*-values were calculated using hypergeometric distribution compared with background peaks and significantly enriched TF motifs were determined with the cutoff *p*-value below 10^-10^. **(c)** From top to bottom, Hi-C contact profiles from untreated (Ctr), Acla- and Daun-treated K562 cells at *Myc* TAD locus. The *Myc* enhancer-promoter loops are indicated by the red dots and enhancer-enhancer loops are indicated by the red rectangles. **(d)** Virtual 4C analysis from the Hi-C data with the viewpoint at the *Myc* gene locus. The interactions between the *Myc* promoter and 7 validated *Myc* enhancers are indicated by red arrows. Normalized CTCF ChIP-seq and virtual 4C signal tracks from untreated (Ctr), Acla- and Daun-treated K562 cells are shown. **(e)** Aggregate peak analysis of CTCF-CTCF loops within ± 50 Kbp window are shown for untreated (Ctr), Acla- and Daun-treated cells. The CTCF-CTCF loops were defined as looping anchors called from Hi-C that overlap with the CTCF peaks from ChIP-seq.

### Daun-induced 3D genome alterations correlate with therapeutic response

Anthracycline-based regimens remain the first-line treatment option for AML due to their efficiency(17). However, patient responses to these therapies are highly variable, influenced by numerous factors(48, 49). To investigate whether the anthracycline-induced alterations in chromatin looping and 3D genome structure contribute to therapeutic response, we performed ATAC-seq on AML blasts isolated from patients and monitored their response to Daun. CD33^+^ or CD34^+^ AML blast cells were isolated via fluorescence-activated cell sorting (FACS) from peripheral blood or bone marrow of 6 complete remission (CR) and 6 refractory/relapsed (R/R) AML patients treated with the DDAG (Daunorubicin+Decitabine+Ara-C+G-CSF) or other anthracycline-based regimens (**Supplementary Table 3**). Samples were first grouped based on the patient response to induction therapy, enabling comparative analysis of Daun-induced chromatin accessibility changes between responders (CR) and non-responders (R/R). Using a distal binarization approach(50), we identified 14,260 group-specific peaks (**Supplementary Table 4**), including 3,357 regions that were closed upon Daun treatment in either CR or R/R patients (**Fig. 6a, Supplementary Fig. 6a**). These closed regions may reflect loss of TF binding and disruption of 3D genome organization, as observed in cell line models (**Fig. 5b**). Motif enrichment analysis on group-specific ATAC-seq peaks revealed that CTCF was significantly affected by Daun treatment between CR and R/R groups (**Fig. 6b**). Similar to K562 cell data, this effect may be attributed to altered CTCF binding rather than differences in CTCF expression or drug uptake, as CTCF levels did not affect K562 viability or correlate with patient survival, and Daun uptake was comparable between the blast cells isolated from the CR and R/R patient groups (**Supplementary Fig. 6b-l**). Notably, the CTCF motif was enriched in regions uniquely closed in CR patients following Daun treatment (**Fig. 6b**), suggesting that additional chromatin loops and 3D genome structures were preferentially disrupted in responders. Pathway enrichment based on the RNA-seq analysis of patient blast cells treated with Daun revealed expected activation of the p53 signaling pathway in both CR and R/R groups (**Supplementary Fig. 6m and n**), along with patient-group-specific pathways, some of which may be potentially linked to 3D genome deregulation (**Supplementary Fig. 6o and p**).

**Figure 6.**
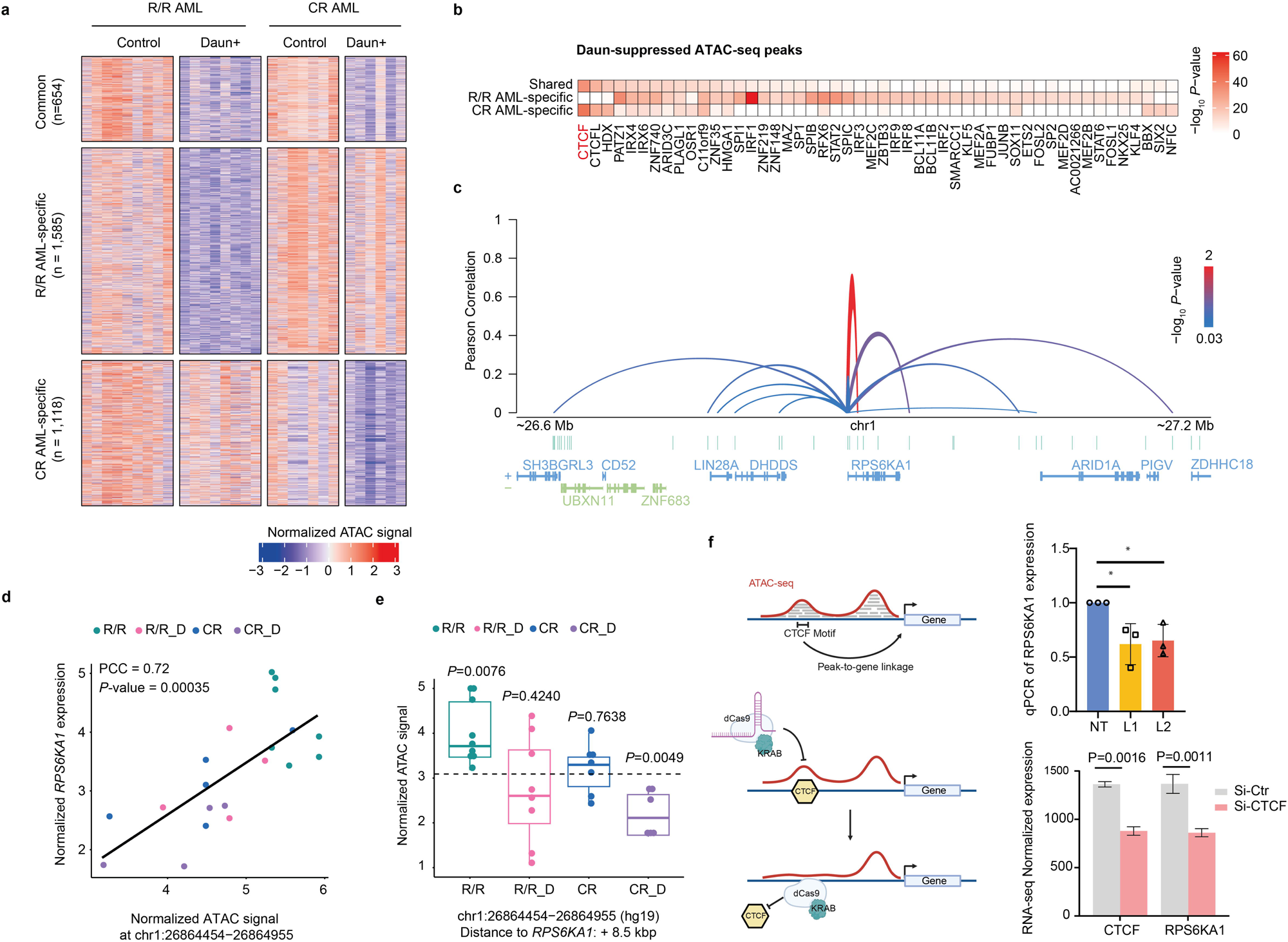
Daun-induced 3D genome alterations in AML correlates with therapeutic response. **(a)** Heatmap of chromatin accessibility in AML patient-specific Daun-suppressed ATAC peaks. AML blasts were treated with 1□µg/mL Daun for 3 hours. CR: complete remission; R/R: refractory/relapsed. Patient-specific Daun-responsive peaks were identified using distal binarization. Differential peaks between groups were assessed by limma eBayes, with FDR < 0.05 defining significance. Significantly suppressed peaks in either the R/R or CR groups are shown. **(b)** TF motif enrichment in patient-subtype-specific Daun-suppressed ATAC peaks. From top to bottom: significantly enriched TF motifs in either shared, R/R AML- or CR AML-specific Daun-suppressed ATAC peak sets are shown. *P*-values were calculated using hypergeometric distribution compared with background peaks and significantly enriched TF motifs were determined with the cutoff *P*-value below 10^-10^. **(c)** Predicted peak-to-gene regulatory links for *RPS6KA1* gene. All ATAC peaks within ±0.5 Mbp of the *RPS6KA1* TSS were tested for correlation between ATAC accessibility and RPS6KA1 expression using paired patient ATAC-seq and RNA-seq data. Pearson correlation coefficients were calculated, and significance was assessed against a null distribution of trans peak-to-gene correlations. Significant links were defined by *P* < 0.05. Arch height of peak-to-gene links represents correlation strength. The color indicates the log_10_ (*P*-value). **(d)** Scatterplot of peak-to-gene links predictive for *RPS6KA1* gene expression, correlating ATAC-seq accessibility with RNA-seq data from individual patients. Pearson correlation coefficient (PCC) and *P*-value are shown. Colors indicate patient-treatment groups; each dot represents one sample. **(e)** ATAC-seq accessibility at CTCF motif-containing peaks across patient groups. The selected peak is predicted to regulate *RPS6KA1* expression based on peak-to-gene link analysis. Signals were normalized to each group’s basal mean accessibility. P-values were calculated using two-tailed Student’s t-test. Colors indicate patient-treatment groups; each dot represents an individual sample. **(f)** Gene expression changes in *RPS6KA1* after perturbation of its cognate CTCF regulatory element.. Left panel: schematic illustrates the strategy of CRISPRi targeting the CTCF-motif-containing ATAC-seq peak to displace CTCF from looping anchor and modulate target gene expression. Right upper panel: qPCR analysis of RPS6KA1 expression in K562 cells after CRISPRi perturbation. NT: cells with dCas9-KRAB and non-targeting RNA; L1/L2: cells with dCas9-KRAB and two distinct guide RNAs targeting the CTCF peak. Error bars show SD of three biological replicates. \**P*<0.05 calculated by two-tailed Student’s t-test. Right lower panel: RNA-seq analysis of *CTCF* and *RPSK6A1* expression after 48h RNAi knockdown of CTCF in K562 cells. si-Ctr: non-targeting siRNA, si-CTCF: siRNA targeting CTCF mRNA. Normalized expression levels of *CTCF* and *RPSK6A1* genes were shown with the standard deviation from 2 biological replicates. Adjusted *P*-values were calculated by DESeq2.

To explore transcriptional consequences of Daun-induced 3D genome changes, we performed peak-to-gene correlation analysis using paired RNA-seq and ATAC-seq of patient blast cells. This analysis, partially recapitulating the physical interactions identified from Hi-C or HiChIP data(50, 51), identified 32,069 peak-to-gene links, including 4,298 involving CTCF (i.e., peaks containing CTCF motifs or overlapping CTCF ChIA-PET loops) (**Supplementary Table 5**). Further stratification based on patient response identified 139 and 211 CTCF-mediated peak-to-gene links uniquely affected by Daun in CR or R/R patients, respectively (**Supplementary Table 6**). To validate predicted regulatory interactions based on peak-to-gene analysis, six CTCF-bound looping regions that may regulate the cell proliferation or drug response-related genes *DNM1*, *DNMT1*, *BAG2*, *RPS6KA1, METTL4,* and *TRIM32,* respectively, were selected for targeting by CRISPR interference (CRISPRi) with dCas9-KRAB(52) (**Fig. 6c-e, Supplementary Fig. 7a-e**). Two links (to *DNM1* and *DNMT1* genes) were specific to R/R patients (**Supplementary Fig. 7a and b**), while four (to *TRIM32*, *BAG2*, *METTL4 and RPS6KA1* genes) were specific to CR patients (**Fig. 6c-e, Supplementary Fig. 7b-d**). CRISPRi targeting of the R/R-specific *DNM1* linkage in K562 cells significantly reduced expression of *DNM1* gene, a GTPase involved in endocytosis (**Supplementary Fig. 7f**)(53). In CR-specific links, CRISPRi targeting the specific peak-to-gene linkages in K562 cells led to significantly reduced expression of *RPS6KA1, TRIM32, BAG2* and *METTL4* genes, respectively (**Fig. 6f, Supplementary Fig. 7f**). RPS6KA1 is a serine/threonine kinase and essential for AML cell survival(54, 55). It has been shown that inhibiting RPS6KA1 sensitizes AML to chemotherapies(56). Therefore, Daun-induced disruption of the *RPS6KA1* regulatory loops may underlie enhanced drug sensitivity in CR patients (**Fig. 6d and e**). We confirmed that *RPS6KA1* downregulation was CTCF-dependent, as shown by RNA-seq analysis of cells with CTCF knockdown using siRNAs (**Fig. 6f**), ruling out the CRISPRi off-target effect(57). Downregulation of *TRIM32* may enhance the p53-mediated apoptosis, cell cycle arrest, and senescence(58), while inhibition of *BAG2* and *METTL4* genes may also contribute to therapeutic response(59, 60). These findings demonstrate that Daun-induced 3D genome alterations may rewire transcriptional regulation via CTCF-mediated long-range interactions in AML patients and contribute to patient-specific treatment responses.

## Discussion

Emerging evidence suggests that aberrant 3D chromatin structures within the human genome contribute to tumor development and malignancy(9, 10). Targeting the 3D genome in cancer cells represents a promising strategy for anti-cancer drug discovery and development. However, current therapeutic options that directly target the 3D genome remain limited. In this study, we identified that anthracycline drugs may redistribute their canonical target, TOP2A, away from the chromatin looping regions. Analyses of TOP2A ChIP-seq and ATAC-seq in Acla- and Daun-treated K562 cells suggest that CTCF binding was disrupted by these drugs. Biochemical assays and CTCF ChIP-seq further confirmed that the chromatin binding of CTCF was disrupted by Acla and Daun treatment. Given the critical role of CTCF in chromatin looping and long-range chromatin interactions, these findings imply that anthracycline drugs may impact 3D genome organization. Indeed, Hi-C analysis revealed that Acla and Daun induced extensive global alterations in long-range chromatin interactions. Moreover, the differential 3D genome targeting between Acla and Daun highlights the potential for personalized use of anthracycline drugs based on patient-specific 3D genome features.

To determine whether similar effects occur in patients, we performed ATAC-seq on blast cells from AML patients treated with Daun. Consistent with our findings in K562 cells, differentially affected ATAC-seq regions in patient blasts were enriched for CTCF motifs, suggesting that CTCF-mediated distal interactions may also be perturbed in patients treated with anthracycline drugs. CRISPRi-mediated targeting of these putative CTCF-bound distal regulatory regions led to reduced expression of linked genes, underscoring their potential therapeutic relevance. Notably, CTCF motifs were enriched in deregulated ATAC-seq regions from Daun-treated CR-patient blast cells, suggesting that 3D genome remodeling may contribute to the enhanced therapeutic response in these patients.

Anthracycline drugs have complex mechanisms of action, including DNA damage and histone eviction(16). Here, we also demonstrate that many anthracycline drugs disrupted 3D genome architecture by targeting CTCF-bound loops and TAD boundaries. While it remains challenging to disentangle the respective contributions of DNA damage, histone eviction, and 3D genome disruption induced by these anthracycline drugs to the overall therapeutic outcome, it is conceivable that tumor cells relying on specific oncogenic 3D genome aberrations may be particularly sensitive to anthracycline-induced 3D genome disruption, such as in AML, where 3D genome rewiring drives malignancy(10). Furthermore, different anthracycline variants exhibited some distinct 3D genome targeting properties, highlighting their potential for personalized therapeutic application. ATAC-seq offers a rapid and informative approach to assess 3D genome responses to anthracycline treatment, with promising clinical relevance for guiding therapy selection. The identification of anthracycline drugs capable of targeting the 3D genome not only offers a new therapeutic strategy for addressing aberrant chromatin architecture in the clinic, but also opens the door for broader exploration of 3D genome dysregulation as an actionable target in cancer and other diseases.

## Materials and Methods

### Primary sample collection

This clinical study was approved by the Human Subject Ethics Committee of Chinese PLA General Hospital. Leukemia blasts were obtained from the peripheral blood or bone marrow of 12 de novo or relapse AML patients with consent (**Supplementary Table 3**). The patients received an intravenous infusion of daunorubicin (60 mg/m^2^) treatment within 3 hours. The AML blast cells were collected before treatment and 3 hours post-daunorubicin administration. The mononuclear cells were isolated from the peripheral blood or bone marrow of donors by density gradient centrifugation using Ficoll-Paque media (GE Healthcare). The CD33+ or CD34+ cells were enriched by magnetic bead (Miltenyi Biotec, Germany) selection.

### Cell culture

K562 cells were cultured in RPMI 1640 + L-Glutamine (Gibco), 10% fetal bovine serum (Biowest), and 1% Penicillin-Streptomycin (Gibco). Cell density and culture conditions were maintained according to the ENCODE Cell Culture Guidelines. K562 cell lines with endogenous TOP2A or CTCF tagged with 3xFLAG were generated as previously described(^61^).

### Reagents

Amrubicin was obtained from Santa Cruz Biotechnology (sc-207289). Aclarubicin was obtained from Santa Cruz Biotechnology (sc-200160). Daunorubicin was obtained from Sanofi-Aventis (the Netherlands). All these drugs were dissolved according to the manufacturer’s formulation, aliquoted and stored at −20 °C for further use.

### Western blot

The K562 cells with CTCF endogenously tagged with FLAG were used for Western blot. 4 million K562 cells were treated with or without 10 μM Daun for 2 hours, followed by washing with cold phosphate-buffered saline (PBS). 1 million cells were lysed in the RIPA buffer (Merck, 20-188) supplemented with protease inhibitor cocktail (Merck,11873580001), and sonicated for 10 cycles of 30s ON and 30s OFF (Bioruptor Pico) at 4 °C. The lysates were centrifuged at 14000 rpm for 15 min at 4 °C, supernatant was collected as the whole cell extract. The remaining 3 million cells were first lysed in hypotonic lysis buffer (50 mM Hepes–KOH, pH 7.5; 140 mM NaCl; 1 mM EDTA; 10% Glycerol; 0.5% NP-40; 0.25% Triton X-100), supplemented with protease inhibitor cocktail and rocked gently at 4°C for 10 mins, centrifuged at 2000 rcf for 4 min at 4 °C. Supernatant was collected as non-chromatin fraction, and the pellets were resuspended in the RIPA buffer (Merck,20-188) supplemented with protease inhibitor cocktail and sonicated for 16 cycles of 30s ON and 30s OFF at 4 °C, vortex briefly for every 4 cycles. Following sonication, samples were centrifuged at 14,000□rpm for 15□min at 4□°C, and the supernatant was collected as the chromatin-bound fraction. Both non-chromatin and chromatin fractions were processed and analyzed by Western blot.

The concentration of protein was determined using the Pierce BCA protein assay kit (Thermofisher). Equal amounts of proteins were run on the 8% SDS-PAGE and were transferred to PVDF membranes. The membranes were blocked with 5% (wt/vol) nonfat milk in PBST for 1 h at room temperature and probed with primary antibodies against Flag (Sigma, F1804) or β-actin (Sigma, A5441), 1:1000 dilution overnight at 4°C. The membranes were washed with PBST three times, and incubated with the appropriate secondary antibodies (1:10000 dilution) for 1 h at room temperature, and then washed with PBST three times. The resulting signal was visualized using the Pierce enhanced chemiluminescence (ECL) Plus Western Blotting Substrate (Thermofisher).

### ATAC-seq

The Omni-ATAC was performed as described previously(62). A detailed procedure and explanation is described in **Supporting Information.**

### ChIP-seq

The ChIP-seq was performed following the previous protocol(63). A detailed procedure and explanation is described in **Supporting Information.**

### In situ Hi-C

In situ Hi-C was performed as previously described(64). A detailed procedure and explanation is described in **Supporting Information.**

### RNA-seq

Total RNA was extracted using TRIzol Reagent (Invitrogen, USA) according to the manufacturer’s instructions. A detailed procedure and explanation is described in **Supporting Information.**

### CRISPRi targeting daunorubicin-responsive peak-to-gene links

The CTCF motif residing in the targeted peaks was identified by FIMO with the cutoff P-value < 10^-5^. The 150□bp motif flanking the genomic sequence was used to identify candidate guide RNA sequences. Two non-overlapping guide RNA sequences of high specificity and targeting either strand were designed per targeted peak-to-gene link using CRISPick (https://portals.broadinstitute.org/gppx/crispick/public). Each guide RNA sequences were individually cloned into the lentiviral vector expressing sgRNA and dCas9-KRAB(65) (Addgene, #71236) following digesting with the BsmBI restriction enzyme. For lentivirus packaging, the 293T cells were grown in 6 well plates. When cells reached 50% confluency, they were transfected with the mixture of 1 µg of Lenti dCas9-KRAB-Puro plasmid, 0.6 µg of psPAX2, 0.4 µg of pCMV-VSV-G, and 6 µg PEI (Polyscience, 23966) in 50 µl of serum-free medium. 8-12 hours post-transfection, the culture medium was refreshed. The media supernatant containing lentiviral particles was collected on the third day after transfection. Then K562 cells were transduced with the virus containing the respective guide RNAs and selected using puromycin (2 µg ml^−1^) for 2-3 days. Expanded cells were collected and verified by qPCR analysis. The sequences of the gRNAs are listed in **Supplementary Table 7**.

### CTCF-overexpressing cell line generation

Single-guide RNAs targeting the promoter regions of the CTCF gene were selected from the pooled CRISPRa screen library(66). The guide RNA sequence was cloned into the lentiSAMv2-Puro plasmid (Addgene, #92062) containing the gRNA scaffold and dCas9 sequence, and lentivirus was made as previously described. Then K562/SAM stable cells (were made by the previous study(14)) were transduced with the virus containing the respective guide RNAs and then selected using puromycin (2 µg ml–1) for 2-3 days. Expanded cells were collected and verified by qPCR analysis. The sequences of the gRNAs are listed in **Supplementary Table 7**.

### RNAi knockdown CTCF

Wildtype K562/K562-TOP2A-FLAG cells were electroporated with small interfering RNA targeting CTCF (Si-CTCF, order from Horizon Discovery) or a non-targeting control siRNA (Si-Ctr, order from Horizon Discovery) using the Lonza 4D-Nucleofector™ system (V4XC-2012) according to the manufacturer’s instructions. Briefly, 1□×□10□ cells were resuspended in electroporation buffer and mixed with 100 pmol siRNA. Transfected cells were immediately transferred to pre-warmed culture medium and incubated for 48-72 hours post-transfection before proceeding with further experiments.

### Circular Chromosome Conformation Capture(4C)-seq

4C-seq was performed as previously described(67). Briefly, 1l710^7^ cells were crosslinked with 2% formaldehyde and quenched by chilled glycine at the final concentration of 0.13 M. Nuclei pellets were isolated by cold lysis buffer (10□mM Tris-HCl pH 7.5, 0.5% NP-40, 1% Triton X-100, 150□mM NaCl, 5□mM EDTA) and supplemented with 1l7 protease inhibitors (Roche). The first digestion was performed overnight at 37□°C with 100U NlaIII enzyme (New England Biolabs). Digestion efficiency was measured by 0.6% agarose gel electrophoresis. DNA was ligated overnight at 16□°C by 50U T4 DNA ligase (Promega) and ligation efficiency was determined by 0.6% agarose gel electrophoresis before reversal of cross-links. The ligated product, also referred to as 3C library, was de-crosslinked by proteinase K and was extracted by phenol-chloroform-isoamyl alcohol mixed solution. The 3C library was then processed overnight at 37□°C for a second digestion with 50U DpnII enzyme (New England Biolabs). After measurement of the second digestion efficiency by 0.6% agarose gel electrophoresis, the second ligation was performed on 25 ug template overnight at 16□°C by 50U T4 DNA ligase (Promega). The ligated product, also referred to as 4C template, was extracted by phenol-chloroform-isoamyl alcohol mixed solution and the concentration was determined by Qubit assays (Thermo Scientific). The 4C template DNA was then amplified by two steps of PCR reactions with Illumina TruSeq adaptors and sent for sequencing on the NovaSeq 6000 platform. Primers used for 4C amplification are listed in the **Supplementary Table 7.**

### Cell viability assay

Cells were seeded in 96-well plates at a density of 10□ cells per well. Amr, Acla or Daun were then added at the indicated concentrations. After 24 hours of treatment, cell viability was assessed using the CellTiter-Blue assay (Promega), following the manufacturer’s instructions.

### FACS analysis of drug uptake

Patient-derived blast cells were treated with 1□µg/ml daunorubicin for 3 hours, and were then analyzed by FACS. The PE-A channel was used to monitor the fluorescence intensity to assess drug uptake as previously described(14).

### Quantitative RT-PCR

Total RNA was isolated from 10^6^ cells using ISOLATE II RNA Mini Kit (Bioline, BIO-52073) according to the manufacturer’s instructions. The cDNAs were synthesized from 600 ng of total RNA using a SuperScript™ IV VILO™ Master Mix (Invitrogen, 11756050) in a 5 μl reaction, and were analyzed using SensiFAST SYBR No-ROX Kit (Bioline, BIO-98020) with gene expression primers for DNM1, DNMT1, BAG2, RPS6KA1, METTL4, TRIM32, and GAPDH (a housekeeping gene) on Biorad CFX Opus 384 Real-Time PCR Systems. The sequences of the primers are in the **Supplementary Table 7**.

### Informatics analysis

Details analysis procedure for ATAC-seq, ChIP-seq, RNA-seq and Hi-C are described in **Supporting Information.**

### Statistics and reproducibility

The western blots for CTCF, and β-actin were repeated three times, and similar results were obtained (**Fig. 3a**). Two biological replications were applied to each ATAC-seq and ChIP-seq experiment in K562 cells. Four biological replicates were applied to each in situ Hi-C experiment in K562 cells. Three biological replications were applied to each qPCR experiment in CRISPRi-editing K562 cells. All other statistics and replications are specified in each method and figure legend. External data used to reproduce the figures is listed in the **Supplementary Table 8**.

## Data availability

The raw sequencing data including ATAC-seq and TOP2A ChIP-seq of cell lines generated in the previous study(61) were deposited at GEO under accession code GSE240443. The raw sequencing data including CTCF ChIP-seq and Hi-C samples of cell lines generated in this study are available at GEO under the accession code GSE276066, GSE285987 (reviewer token: gfklgaumjvetfaf). All the processed ChIP-seq and ATAC-seq bigwig signal files including primary AML samples are available under the UCSC genome browser session (https://genome-euro.ucsc.edu/s/mtan995/hg19_ANTH_AML).

## Code availability

The scripts for reproducing the genomic data analysis are available on GitHub at (https://github.com/MorganTan95/ANTH_3D_genome_paper).

## Acknowledgements

This work was supported by the Gisela Thier Fellowship from Leiden University Medical Center and the ERC Starting Grant 950655-Silencer from the European Research Council (all awarded to B.P.), and by the National Institutes of Health (R35-GM128645 to D.H.P. and 5T32 CA217824 to A.A.P.).

## Author contribution

B.P. conceptualized the project. M.T., S.S., Y.L., A.A.P., and B.P. conducted investigations. M.T., S.S., A.A.P., and D.H.P conducted formal analysis. M.T., S.S., and B.P. wrote the original draft. B.P., M.T., S.S., and D.H.P. reviewed and edited the manuscript. B.P., D.L., and D.H.P. supervised the project.

## Declarations of Interests

The authors declare no competing interests.

## Notes

### Competing Interest Statement

The authors have declared no competing interest.

### Summary of Updates

1. the main figures is uploaded

