## Supporting Information for "Targeting the 3D genome by anthracyclines for chemotherapeutic effects"

**Supporting Information of Materials and Methods**

**ATAC-seq**

The Omni-ATAC was performed as described previously(1). Briefly, the 50,000 wildtype K562 cells were treated with Amr, Acla or Daun at a concentration of 10 µM for 2 hours. Cells were then harvested and washed with 1 ml cold resuspension buffer (RSB, 10 mM pH 7.4 Tris-HCl +10 mM NaCl + 3 mM MgCl2 + ddH_2_O), and lysis in 50 µl cold ATAC lysis buffer (RSB + 0.1% v/v Tween + 0.1% v/v NP40 + 0.01% v/v Digitonin) on ice for 3 mins to isolate nuclei. The lysed cells were then washed in RSB buffer by gently inverting the tube three times. Nuclei were pelleted by centrifugation at 500 rcf for 10 mins at 4 °C. The supernatant was carefully discarded to avoid disturbing the nuclear pellet. Nuclei pellets were resuspended in a 50 µl transposition reaction mix (2x TD buffer, 2.5 µl transposase (100 nM final), 16.5 µl PBS, 0.5 µl 1% digitonin, 0.5 µl 10% Tween-20, 5 µl nuclease-free water). The mixture was triturated 6 times and incubated at 37°C for 30 mins on a thermomixer set to 1000 rpm. Transposed DNA fragments were purified using Zymo DNA Clean and Concentrator-5 kit (Cat. No. D4014) and subsequently amplified by PCR. The PCR reaction consisted of 20 µl purified transposed DNA, 2.5 µl 25 µM PCR primer 1 (AD 1), 2.5 µl 25 µM Barcoded PCR primer 2 (AD 2.1 - AD 2.5), and 25 µl NEB Next High-Fidelity 2x PCR Master. The thermal cycle is as following: 1 cycle: 5 min 72°C, 30 sec 98°C; 5 cycles: 10 sec 98°C, 30 sec 63°C, 1 min 72°C. The concentration of pre-amplified fragments was determined by quantitative PCR (qPCR), and this concentration was used to estimate the total number of cycles required to obtain fragments. QPCR reaction was set up as 5 µl DNA from the last PCR step, 1.6 µl nuclease-free H_2_O,1.25 µl 5 µM PCR primer 1 (AD1), 1.25 µl 5 µM Barcoded PCR primer 2 (AD2.1 - AD 2.5), 0.9 µl 10x SYBR Green I, and 5 µl NEB Next High-Fidelity 2x PCR Master Mix. Using a qPCR instrument, cycle as follows: 1 cycle: 30 sec 98°C, 20 cycles: 10 sec 98°C, 30 sec 63°C, 1 min 72°C. To determine the additional number of PCR cycles needed, plot the linear Rn (normalized fluorescence) versus cycle number. Identify the cycle number corresponding to one-third of the maximum fluorescence intensity; this value should be less than 7 and will be used to define the optimal number of additional amplification cycles. A second PCR was performed to amplify the pre-amplified fragments for the desired number of cycles. Second PCR fragments were purified using Zymo DNA Clean and Concentrator-5 kit and were run on the E-gel. DNA fragments ranging from 150 bp to 800 bp were size selected for downstream analysis. The excised gel fragment was purified using the MinElute Gel Extraction Kit (QIAGEN, Cat. No. 28606) and eluted in 15 µL of nuclease-free water. DNA concentration (ng/µL) was measured using a Qubit fluorometer, and the quality of the purified libraries was assessed using the Agilent Bioanalyzer with the High Sensitivity DNA Analysis Kit.

**ChIP-seq**

The ChIP-seq was performed following the previous protocol(2). All lysis buffers should be supplemented with protease inhibitors (Complete, EDTA-free, Roche, #11873580001). The 10^8^ wildtype K562 cells and K562-TOP2A-3xFLAG tag cells were treated with Amr, Acla or Daun at a concentration of 10 µM for 2 hours, as indicated. Following treatment, the cells were crosslinked with 1% formaldehyde and incubated for 10 mins at room temperature with constant gentle stirring to ensure efficient cross-linking. The crosslinking reaction was quenched by adding 2 M glycine to a final concentration of 125 mM, followed by incubation for 5 minutes at room temperature. Cells were then washed with PBS and pelleted by centrifugation. The cell pellet was lysed in 10 ml of lysis buffer 1 (50 mM Hepes-KOH, pH 7.5; 140 mM NaCl; 1 mM EDTA; 10% Glycerol; 0.5% NP-40 or Igepal CA-630; 0.25% Triton X-100) at 4°C for 10 mins. Nuclei were pelleted by centrifugation at 2000 rcf for 4 mins at 4°C and resuspended in 10 ml of lysis buffer 2 (10 mM Tris–HCl, pH 8.0; 200 mM NaCl; 1 mM EDTA; 0.5 mM EGTA) and rocked gently at 4 °C for 5 mins. Chromatin was pelleted by centrifugation at 2000 rcf for 4 mins at 4°C, resuspended in 2 ml of 1XRIPA buffer (Merck, 20-188) with washed sonication beads, and sheared with a Bioruptor® Pico sonication device. The Chromatin pellet should be kept in an ice-water bath during sonication at 4°C, and sonication using 16 cycles of 30s ON and 30s OFF (every 4 cycles, stop the sonication and vortex briefly). The supernatant was collected by centrifugation at 3000 rpm for 5 mins at 4°C and mixed with Triton X-100 to make the final concentration of Triton X-100 to 1% in 1.5 ml tubes. Insoluble chromatin and debris were removed by centrifugation at 14,000 rpm for 15 mins at 4°C.

The supernatant was incubated with 100 µl antibody/magnetic bead mix. Magnetic beads should be pre-blocked with antibody against CTCF (Merck. 07-729) or Flag (Sigma, F1804) overnight or for a minimum of 4 h on a rotating platform at 4°C. The antibody-conjugated magnetic beads were washed three times with 1 ml of blocking solution (0.5% BSA [w/v] in PBS) and resuspended in 100 µl of the same blocking solution in a 1.5 mL tube. The beads were then incubated overnight at 4 °C on a rotator or rocker. Bound fragments- antibody/magnetic beads were washed with 1 ml RIPA Buffer for 4-6 more times, then washed once with 1 ml TBS (20 mM Tris–HCl, pH 7.6; 150 mM NaCl). Followed by centrifugation at 960 rcf for 3 mins at 4°C and placement on a magnetic stand to remove any residual TBS buffer. Bound fragments-antibody/magnetic beads were resuspended in 200 µl of elution buffer (50 mM Tris–HCl, pH 8; 10 mM EDTA; 1% SDS) for reverse crosslinking at 65 °C for 6-18 h on a thermomixer with 1000 rpm (50 µl of the WCE was reversed the formaldehyde crosslinking simultaneously with the ChIP samples). The supernatant was transferred to a new tube and incubated with 200 µL of TE buffer to dilute the SDS present in the elution buffer, along with 8 µL of RNase A (1 mg/mL; Ambion, Cat. No. 2271) at 37 °C for 30 minutes. The sample was then incubated with 4 µl of 20 mg/ml proteinase K (Invitrogen, 25530-049) at 55°C for 1-2 h. The ChIP DNA was cleaned up using QIAquick PCR Purification Kit (QIAGEN, 28106) and eluted in 35 µl EB.

A total of 34 µl ChIP DNA was subjected to end-repair in a 50 µL (Lucigen,14422-5). Incubated at room temperature for 45 mins and purified using the QIAquick PCR Purification Kit (QIAGEN, 28106), eluting in 35 µl of EB. Then the 34 µl end-repaired ChIP DNA was added of ‘A’ base to 3’ Ends in a 50 µl reaction, with 5 µl Klenow buffer (NEB2), 10 µl 1 mM dATP, 1 µl Klenow (3’ to 5’ exo minus). Incubated for 30 mins at 37°C and purified using the QIAquick PCR Purification Kit (QIAGEN, 28106) eluting in 12 µl of EB. The 12 µl purified DNA was ligated with an adapter in a 30 µl reaction, with 15 µl 2x DNA ligase buffer,1 µl Adapter oligo mix (40 ng/µl), and 2 µl T4 DNA ligase. Incubated at room temperature for 15 minutes and purified using the QIAquick PCR Purification Kit (QIAGEN, 28106), eluting in 25 µl of EB. The 23 µl purified DNA was combined and mixed with 25 μl Phusion High-Fidelity PCR Master Mix (Thermofisher, F531L), 1 µl 10 µM PCR primer 1.1, and 1 µl 10 µM PCR primer 2.1 to a total 50 µl reaction volume. PCR procedures were 98 °C for 30 s, 15 cycles of 98 °C for 10 s, 65 °C for 30 s, 72 °C for 30 s, and 72 °C for 5 min (15 cycles could be used for high abundant ChIP DNA, 17 cycles could be used for low abundant ChIP DNA). The PCR product was purified using the QIAquick PCR Purification Kit (QIAGEN, 28106), and eluted in 19 µl of EB. The adapter-ligated DNA was separated on a 2% Agarose E-gel (ThermoFisher, G402022) and the most abundant approximately 300 bp DNA band (and within the expected size range of 450–600 bp) was excised from the gel using a clean scalpel. Gel images were documented before and after excision to ensure accurate recovery. The excised gel fragment was purified using MinElute Gel Extraction Kit (QIAGEN, 28606), and eluted in 15 µl of EB buffer. DNA concentration (ng/µl) was measured using a Qubit fluorometer, and library quality was assessed with the Agilent Bioanalyzer using the High Sensitivity DNA Analysis Kit.

**In situ Hi-C**

In situ Hi-C was performed as previously described(3). The 10^7^ wildtype K562 cells were treated with Acla or Daun at a concentration of 10 µM for 2 hours. The cell pellet was lysed in ice-cold Hi-C lysis buffer (10 mM Tris-HCl pH 8.0, 10 mM NaCl, 0.2% IGEPAL CA630) with 50 μl of protease inhibitors for 15 mins on ice. The lysed cells were then pelleted and washed once using the ice-cold Hi-C lysis buffer, and resuspended in 50 μl of 0.5% SDS and incubated at 62°C for 7 mins. Reactions were quenched with 145 μl water and 25 μl 10% Triton X-100 at 37°C for 15 mins. Chromatin was digested overnight with 25 μl of 10X NEBuffer 2 and 100 U of MboI at 37°C with rotation. The MboI enzyme was inactivated by incubation at 62 °C for 20 minutes, followed by cooling to room temperature. Fragment overhangs were repaired by adding 37.5 μl 0.4 mM biotin-14-dATP, 1.5 μl each 10 mM dCTP, dGTP, dTTP, 8 μl 5 U/μl DNA Polymerase I, Large (Klenow) fragment and incubating at 37°C for 1.5 h with rotation. Ligation was performed by adding 673 μl water, 120 μl 10x NEB T4 DNA ligase buffer, 100 μl 10% Triton X-100, 6 μl 20 mg/mL BSA, and 1 μl 2000 U/μl T4 DNA ligase and incubating at RT for 4 h with slow rotation. The ligated samples were pelleted by centrifugation at 2,500 × g and resuspended in 432 μl of nuclease-free water containing 18 μl of proteinase K (20 mg/mL), 50 μl of 10% SDS, and 46 μl of 5 M NaCl. The mixture was incubated at 55 °C for 30 minutes, followed by overnight incubation at 68 °C. The samples were cooled to room temperature, then mixed with 1.6× volumes of 100% ethanol and 0.1× volumes of 3 M sodium acetate (pH 5.2), followed by incubation at −80 °C for 4–6 hours. The samples were then centrifuged at maximum speed (≥13,000 × g) at 2 °C for 15 minutes and washed twice with 70% ethanol. The resulting DNA pellet was dissolved in 130 μl of 10 mM Tris-HCl (pH 8.0), incubated at 37 °C for 1–2 hours, and stored at 4 °C overnight. The resulting DNA was sheared using the Covaris LE220 (Covaris, Woburn, MA) to a fragment size of 300-500 bp in a Covaris microTUBE. The sheared DNA was transferred to a fresh tube and the Covaris microTUBE was rinsed with 70 μl of water and pooled with the sample. A 1:5 dilution of the sheared DNA was run on a 2% agarose gel to verify fragment size distribution. Size selection was then performed using AMPure XP beads. The cleaned-up undiluted DNA was run on a 2% agarose gel to verify size selection between 300-500 bp. A total of 150 μl of 10 mg/mL Dynabeads MyOne Streptavidin T1 beads were washed with 400 μl of 1× Tween Washing Buffer (TWB; composed of 250 μl Tris-HCl [pH 7.5], 50 μl 0.5 M EDTA, 10 mL 5 M NaCl, 25 μl Tween-20, and 39.675 μl nuclease-free water). The beads were then resuspended in 300 μl of 2× Binding Buffer (500 μl Tris-HCl [pH 7.5], 100 μl 0.5 M EDTA, 20 ml 5 M NaCl, and 29.4 ml nuclease-free water), added to the DNA sample, and incubated at room temperature for 15 minutes with rotation. DNA-bound beads were then washed twice with 600 μl of 1x TWB at 55°C for 2 mins with shaking and resuspended in 100 μl 1x NEBuffer T4 DNA ligase buffer, transferred to a new tube, and reclaimed. Sheared ends were repaired by resuspending the beads in 88 μl of 1x NEB T4 DNA Ligase Buffer with 1mM ATP, 2 μl of 25 mM dNTP mix, 5 μl of 10 U/μl NEB T4 PNK, 4 μl of 3 U/μl NEB T4 DNA polymerase I, and 1μl of 5U/μl NEB DNA polymerase 1, large (Klenow) fragment and incubating at RT for 30 mins. The beads were washed twice with 1× TWB for 2 minutes at 55 °C with shaking, followed by a single wash with 100 μl of 1× NEBuffer 2. The beads were then transferred to a new tube and resuspended in a reaction mixture containing 90 μl of 1× NEBuffer 2, 5 μL of 10 mM dATP, and 5 μl of NEB Klenow Fragment (exo–). The mixture was incubated at 37 °C for 30 minutes. The beads were then washed twice with 1× TWB for 2 minutes at 55 °C with shaking, followed by a wash with 100 μl of 1× Quick Ligation Reaction Buffer. The beads were subsequently transferred to a new tube and reclaimed using a magnetic stand. The beads were resuspended in 50 μl of 1× NEB Quick Ligation Reaction Buffer containing 2 μl of NEB Quick DNA Ligase and 3 μl of an appropriate Illumina indexed adapter (TruSeq Nano), and incubated at room temperature for 15 minutes. After ligation, the beads were reclaimed using a magnetic stand and washed twice with 1× TWB for 2 minutes at 55 °C. They were then washed once with 100 μl of 10 mM Tris-HCl (pH 8.0), transferred to a new tube, reclaimed again, and finally resuspended in 50 μl of 10 mM Tris-HCl (pH 8.0).

Hi-C libraries were amplified directly from the T1 beads using 8 PCR cycles. The reaction consisted of 5 μl of PCR primer cocktail, 20 μl of Enhanced PCR Mix, and 25 μl of bead-bound DNA. PCR conditions were as follows: initial denaturation at 95 °C for 3 minutes, followed by 4-12 cycles of 20 seconds at 98 °C, 15 seconds at 60 °C, and 30 seconds at 72 °C. Following amplification, samples were held at 72 °C for 5 minutes for final extension, then cooled and held at 4 °C. Amplified samples were purified using AMPure XP beads. The library concentration was measured using a Qubit fluorometer, and library quality was assessed with the Agilent Bioanalyzer using the High Sensitivity DNA Analysis Kit. The resulting libraries were subjected to paired-end 2 × 150 bp sequencing on an Illumina NovaSeq platform.

**Primary AML samples ATAC-seq library construction**

The Hyperactive ATAC-Seq Library Prep Kit for Illumina (Nanjing Vazyme Biotech Co, TD711-01) was used for AML sample ATAC-seq library construction. Briefly, cells were gently thawed at RT to maximize the cell viability recovery and counted. For each sample library preparation, 10^5^ cells were pelleted by centrifugation and washed twice with 50 μl of pre-cooled Tagment DNA & Wash Buffer. 10⁵ cells were treated with 1 µg/mL daunorubicin (Daun) for 3 hours before harvesting for library preparation. The cell pellet was resuspended in 50 µL of pre-cooled lysis buffer (48.5 µL RS Buffer, 0.5 µL 10% NP-40, 0.5 µL 10% Tween-20, and 0.5 µL 1% digitonin) and incubated on ice for 5 minutes. The cells were centrifuged at 2,300 rpm (500 × g) for 10 minutes at 4 °C, and the supernatant was gently removed. Then, the cells were resuspended in 50 μl of precooled fragmentation mix (16.5 μl TW Buffer, 0.5 μl 10% Tween-20, 0.5 μl 1% Digitonin, 18.5 μl Nuclease-free ddH2O, 10 μl 5 × TTBL and 4 μl TTE Mix V50). The mixture was gently pipetted up and down 10 times to ensure thorough mixing, followed by incubation in a water bath at 37 °C for 30 minutes. After the tagmentation reaction, the mixture was gently pipetted to mix thoroughly and transferred to a 200 μl PCR tube. Subsequently, 5 μl of Stop Buffer was added to each tube, the cells were resuspended, and the samples were incubated at room temperature for 5 minutes. To extract the fragmented DNA, 100 μl of well-mixed ATAC DNA Extract Beads was added to 55 μl of the terminated sample. The mixture was gently pipetted to mix and incubated at room temperature for 5 minutes. The magnetic beads were washed twice with 200 μl of freshly prepared 80% ethanol, incubated at room temperature for 30 seconds during each wash. After each wash, the supernatant was carefully removed without disturbing the beads. After air-drying the beads for 5 minutes, the reaction tube was removed from the magnetic rack, and the DNA was eluted with 26 μl of nuclease-free water. The mixture was incubated at room temperature for 5 minutes to ensure complete elution. Centrifuge the reaction tube briefly and place it on the magnetic rack to separate the magnetic beads from the solution. Wait 5 min until the solution becomes clear, then carefully pipette 24 μl of the supernatant into a new PCR tube. The library was amplified using 60 ul PCR reaction as 20 μl purified fragmented DNA, 30 μl 2×CAM, 5 μl N5XX PCR primer, and 5 μl N7XX PCR primer. The PCR program is as follows: 3 min 72°C, 3 min 95°C; 12 cycles of 10 sec 98°C and 5 sec 60°C, 1 min 72°C. The PCR product was size selected using ATAC DNA Clean Beads by mixing 33 μl (0.55 ×) magnetic beads into 60 μl PCR products gently followed by incubation at RT for 5 min. Wash the magnetic beads twice with 200 μl of freshly prepared 80% ethanol, incubate at RT for 30 sec, and carefully remove the supernatant. After air-drying for 5 min, remove the reaction tube from the magnetic rack and elute with 22 μl of Nuclease-free ddH2O followed by incubation at RT for 5 min. Centrifuge the reaction tube briefly and place it on the magnetic rack to separate the magnetic beads from the solution. Wait 5 min until the solution becomes clear, then carefully pipette 20 μl of the supernatant into a new sterile tube. The library concentration was measured using a Qubit fluorometer, and library quality was assessed with the Agilent Bioanalyzer using the High Sensitivity DNA Analysis Kit.

**Primary AML samples RNA library construction and sequencing**

10⁵ cells were treated with 1 µg/ml daunorubicin (Daun) for 3 hours before harvesting for library preparation. mRNA was then purified from 5 µg of total RNA using Dynabeads Oligo(dT) (Thermo Fisher, CA, USA) with two rounds of purification to ensure high specificity and yield. mRNA fragmentation was performed using divalent cations with the Magnesium RNA Fragmentation Module (NEB, Cat. No. E6150, USA) by incubating at 94 °C for 5–7 minutes. The fragmented RNA was then reverse-transcribed into cDNA using SuperScript™ II Reverse Transcriptase (Invitrogen, Cat. No. 1896649, USA). The resulting first-strand cDNA was subsequently used to synthesize U-labeled second-strand DNA using E. coli DNA Polymerase I (NEB, Cat. No. M0209, USA), RNase H (NEB, Cat. No. M0297, USA), and dUTP Solution (Thermo Fisher, Cat. No. R0133, USA). An A-base was then added to the blunt ends of each strand, preparing them for ligation to the indexed adapters. Each adapter contained a T-base overhang for ligating the adapter to the A-tailed fragmented DNA. Dual-index adapters were ligated to the fragments, and size selection was performed with AMPureXP beads. Following treatment of the U-labeled second-strand DNA with heat-labile UDG enzyme (NEB, Cat. No. M0280, USA), the ligated products were amplified by PCR under the following conditions: initial denaturation at 95℃ for 3 min; 8 cycles of denaturation at 98℃ for 15 sec, annealing at 60℃ for 15 sec, and extension at 72℃ for 30 sec; and then final extension at 72℃ for 5 min. The final cDNA libraries had an average insert size of 300 ± 50 bp. Paired-end sequencing (2 × 150 bp, PE150) was performed on an Illumina NovaSeq™ 6000 platform (LC-Bio Technology Co., Ltd., Hangzhou, China) according to the manufacturer’s recommended protocol.

**ATAC-seq data processing**

The ATAC-seq samples from K562 cells were processed using the ENCODE Data Coordination Center (DCC) ATAC-seq pipeline (v1.10.0) (<https://github.com/ENCODE-DCC/atac-seq-pipeline>). Briefly, the ATAC-seq reads were aligned to the human reference genome (GRCh37/hg19) using Bowtie2(4). Duplicated and mitochondrial reads were filtered. The ATAC-seq fragment sizes for all the samples were corrected for the Tn5 offset (“+” stranded +4 bp, “-” stranded -5 bp) and plotted using the geom_density function in ggplot2.

**Enrichment analysis of Tn5 insertion sizes within chromatin states**

The ATAC-seq insertion size enrichment analysis in different chromatin states was performed as described(5). First, the Tn5 insertion sizes ranging from 50 bp to 750 bp were divided into 100 bins and the percentage of insertions in each bin was calculated, followed by normalizing to the maximal percent within each state. Finally, the enrichment of different Tn5 insertion sizes within each state was calculated by normalizing to the genome-wide different Tn5 fragment sizes.

**Nucleosome positioning analysis**

To define the regions of nucleosome analysis, broad peaks were first called from all the ATAC-seq samples identically using MACS2(6) with the settings of ‘-p 0.05 --broad --broad-cutoff 0.05 --keep-dup all’ and then extended by 200 bp on either sides. The python package NucleoATAC(7) (v0.3.4) was used to call nucleosome positions and occupancy within the defined regions from low-quality reads filtered and non-mitochondrial BAM files. The predicted nucleosome occupancy scores signal tracks from different samples by NucleoATAC around interesting regions were compared and plotted by deepTools(8).

**TF footprinting analysis**

The TF footprints were identified using HINT-ATAC(9) within the broad peaks called by MACS2 with the settings of ‘-p 0.05 --broad --broad-cutoff 0.05 --keep-dup all’. The TF motifs from JASPAR database(10) were used to match within the predicted footprints to infer the bound (active) TFs. The average Tn5 cleavage profiles around binding sites of particular TFs were plotted and the TF activity scores were calculated to compare TF binding strength under different drug treatment conditions.

**Differential chromatin accessibility analysis**

The R package csaw(11) were used for differential chromatin accessibility analysis as suggested(12). Briefly, filtered BAM files of two biological replicates from two conditions are specified as the inputs for the csaw pipeline. The genome were segmented into 300bp windows and reads count within each window were computed for each sample. The windows with low reads enrichment (< 3-fold threshold relative to 3 Kbp neighborhood) were filtered. TMM normalization was implemented for read counts within each window and de novo locally enriched windows were used as the query regions for differential accessibility comparison. Differentially accessible nearby windows within 100 bp apart were merged. Significant differential regions were filtered using the threshold of FDR < 0.05.

**ChIP-seq data processing and visualization**

The ChIP-seq data were processed using the ENCODE Data Coordination Center (DCC) ChIP-seq pipeline (<https://github.com/ENCODE-DCC/chip-seq-pipeline2>) (v1.9.0). Briefly, the ChIP-seq reads were aligned to the human reference genome (GRCh37/hg19) using Bowtie2(4). Duplicate reads were removed using Picard MarkDuplicates (https://broadinstitute.github.io/picard/). Peaks of each biological replicate were called against the whole-cell lysates (WCL) replicates using SPP(13) with the parameters ‘-npeak 300000 -speak 155 -fdr 0.01’. Reproducible peaks were intersected from two biological replicates. The blacklisted regions described by ENCODE were discarded.

The ChIP-seq signal tracks were generated from BAM files using the command ‘bedtools genomecov -scale’ with read count per million (CPM) normalization and converted to bigwig files using bedGraphToBigWig(14). The genome snapshots were created with  Integrative Genomics Viewer (IGV) browser(15). The heatmap of ChIP-seq signal in the regions of interest and similarity comparisons of genome-wide ChIP-seq signal distribution were computed by deepTools(8).

**Differential ChIP-seq peaks calling**

The R package csaw(11) were used for differential peak calling for TOP2A and CTCF ChIP-seq experiments. Briefly, filtered BAM files of two biological replicates from two conditions are specified as the inputs for the csaw pipeline. The genome was segmented into 20bp windows and reads count within each window were computed for each sample. The windows with low reads enrichment (< 3-fold threshold relative to 2Kb neighborhood) were filtered. TMM normalization was implemented for read counts within each window and de novo locally enriched windows were used as the query regions for differential binding affinity comparison. Differential nearby windows within 50 bp apart were merged. Significant differential ChIP-seq peaks were filtered using the threshold of FDR < 0.05. Motif enrichment analysis within the differential ChIP-seq peaks was performed by HOMER(16).

**Permutation association test for ChIP-seq peak sets**

The associations between two ChIP-seq peak sets were determined by calculating the significance of overlapping peaks based on the genome coordinates. The function enrichPeakOverlap() in R package ChIPseeker(17) were used to shuffle the genome coordinates of the target peak set and calculate the overlaps of the query peak set with the shuffled target peak set. The permutation step was repeated for 50,000 times to estimate the null distribution. Then the P-values were calculated and adjusted by the Benjamini-Hochberg (BH) method.

**Hi-C data processing and visualization**

In situ Hi-C datasets were processed using a modified version of the Juicer Hi-C pipeline (<https://github.com/EricSDavis/dietJuicer>) with default parameters as previously described(18). Reads were aligned to the GRCh37/hg19 human reference genome with bwa (v0.7.17) and MboI was used as the restriction enzyme. Four biological replicates were aligned and merged. Chromatin loops were called using the supervised machine learning framework called Peakachu(19) which has been pre-trained at different sequencing depth to overcome bias from different samples. The replicates-merged, MAPQ > 30 filtered and Knight & Ruiz methods (KR) balanced .hic matrices were used as inputs for calling loops at 10Kbp resolution with respective sequencing depth models (control: 1.2 billion model, Acla: 900 million model, Daun: 850 million model) and the cut-off of loop probability above 0.9. This resulted in 37,479 loops for control, 15,347 loops for Acla-treated and 9,438 loops for Daun-treated samples after filtering.

For visualizing the Hi-C matrices, the R package plotgardener(20) was used with the KR-normalized Hi-C matrices as input. To get the idea of global chromatin folding pattern change after anthracycline treatment, we plotted the *P(s)* curves of samples either with or without anthracycline treatment using the cooltools workflow(21) (https://github.com/open2c/open2c_examples/blob/master/contacts_vs_distance.ipynb), which compared the rate of decay of contacts with the genomic separation distance across different samples.

**TAD boundaries calling and visualization**

To identify the TAD boundaries, we use the diamond insulation algorithm(22) implemented in cooltools pipeline(21) with the input of KR-balanced Hi-C matrix and the parameters setting of 10 kb bin size and 100 kb window size. The genome-wide boundary scores (*BS*) and insulation scores (*IS*) were calculated as output. The 10 kb bins satisfying the cutoff of 0.2 ≤ *IS* < 0.5 and *IS* ≥ 0.5 were called as weak and strong boundaries respectively as suggested (https://data.4dnucleome.org/resources/data-analysis/insulation_compartment_scores). TAD boundaries from published Hi-C were defined as ±25 kb windows centered at TAD end positions(23).

The differential boundary analysis was performed as previously described(24). Briefly, lost boundaries by anthracycline treatments were defined by the diamond insulation algorithm, i.e. same locus bins not called as boundaries in drug-treated samples but called as boundaries in the control sample. Residual boundaries by anthracycline treatments were defined if the locus bins called as boundaries in drug-treated samples. No gained boundaries were reported in Acla- and Daun-treated samples by the diamond insulation pipeline. The bins with *BS^control^ - BS^drug^* ≥ 0.4 and *IS^control^ - IS^drug^* ≤ -0.1 were defined as weakened or less insulated boundaries by drug treatment. The bins with *BS^control^ - BS^drug^* ≤ -0.4 and *IS^control^ - IS^drug^* ≥ 0.1 were defined as weakened or less insulated boundaries by drug treatment. The other residual boundaries were regarded as stable.

The correlation of boundary scores and insulation scores of residual TAD boundaries after anthracycline treatment were plotted using the R package ggplot2. The local pileups of lost or differential boundaries were calculated using coolpup.py(25) with the parameters ‘--local --ignore_diags 0 --view --flank 500000’ and visualized with plotpup.py.

**Differential chromatin loops calling**

To identify the drug-induced lost and gained loops, we relied on the genome-wide loop probability prediction by Peakachu(19), which is positively associated with loop intensity and the fold-change of Peakachu probability at the same locus in two individual samples can be used to measure the looping dynamics(26) (i.e. gained or lost). A Gaussian mixture model was used to fit the reciprocal fold-changes of the Peakachu probabilities in the merged loop regions which was called either from control or anthracycline-treated samples. The threshold of FDR below 0.05 was used as the threshold to extract the significant differential loops. This procedure resulted in 6,328 gained and 24,460 lost loops by Acla treatment and 4,800 gained and 28,025 lost loops by Daun treatment. The non-overlapped loops were regarded as stable loops.

**Aggregate peak analysis**

To assess the quality of differential loops calling, we generated the aggregate peak analysis (APA) plots and scores by using the *apa-analysis* script in HiCPeaks (<https://github.com/XiaoTaoWang/HiCPeaks/>) with the default parameters. Briefly, the pixels within 5 bins in both directions were extracted (11x11) and the normalized contact signals at each position were calculated by dividing the observed contacts by the expected contacts at the distance accordingly. The mean of observed over expected ratios at each position of the matrix was assigned to the pixel and plotted out. The APA score was calculated by dividing the center pixel value by the mean value of the nine pixels (3x3) in the lower right section. The z-score was calculated by dividing the difference between center pixel value and mean of the lower right section by the standard deviation of the lower right section. The p-value was converted from z-score using one-side t-test.

The APA analysis around *cis*-regulatory loops were performed as described above. To determine different types of *cis*-regulatory loops, the CTCF-CTCF loop was called by overlapping the both anchors of Hi-C loop with CTCF ChIP-seq peaks, the promoter-promoter loop was called  by overlapping both anchors of Hi-C loop with the promoter regions (within -1500 to +500 bp from TSS) presenting active promoter mark (H3K4me3 peaks), the enhancer-promoter loops by overlapping one anchor of Hi-C loop with distal regions (excluding promoters) presenting active promoter mark (H3K27ac peaks), the other anchor with active H3K4me3 promoters, the enhancer-enhancer loops by overlapping both anchors of Hi-C loops with the active H3K27ac distal enhancers.

**Virtual 4C analysis**

The track for the virtual 4C analysis anchored at *Myc* promoter or 3’ HS1 site based on Hi-C data was generated using CoolBox(27) with default parameters. The 5 kb resolution Hi-C matrix was used as input and the validated 7 *Myc* enhancers(28) or HS5 LCR site(29) were highlighted.

**KEGG pathway enrichment in differential chromatin loops**

KEGG pathway enrichment was performed on the genes whose promoters (within -1500 to +500 bp from the TSSs) overlapped either a gained or lost looping anchor after the drug treatments using clusterProfiler(30) (v4.12). KEGG pathway terms were filtered using the cutoff of adjusted p-value below 0.05.

**AML RNA-seq data processing**

The adaptors were trimmed from paired-end RNA-seq reads using Cutadapt (v1.9). The trimmed reads were aligned to the human reference genome (GRCh37/hg19) using HISAT2(31) (v2.2.1). The mapped reads were assembled using StringTie(32) (v2.1.6) with default parameters. Then, all transcriptomes from all samples were collapsed to reconstruct a comprehensive transcriptome using gffcompare(33) (v0.9.8). After the final transcriptome was generated, StringTie and ballgown(34) were used to estimate the expression levels of all transcripts and perform expression abundance for mRNAs by calculating FPKM (fragment per kilobase of transcript per million mapped reads) value.

**AML ATAC-seq data processing**

The ATAC-seq samples from primary AML patients were identically processed using the Snakemake pipeline (https://github.com/porchard/ATACseq-Snakemake). Briefly, the ATAC-seq adapters were trimmed from the paired-end ATAC-seq reads using cta (v0.1.2) (https://github.com/ParkerLab/cta). The trimmed reads were aligned to the human reference genome (GRCh37/hg19) using BWA-MEM. The duplicated reads were removed using Picard MarkDuplicates. The low-quality mapped and mitochondrial reads were filtered.

**Reproducible ATAC-seq peaks calling from AML samples**

Peaks were called using MACS2 with the parameters of ‘-p 0.01 --shift -75 --extsize 150 --nomodel --call-summits --nolambda --keep-dup all’, then followed by blacklisted regions filtering recommended by ENCODE(35) and peaks mapped to chromosome Y removal. To create a single merged ATAC-seq peak set for all samples, we first categorize all the samples based on a) the patient's response assessment of daunorubicin-based chemotherapy: complete response (CR) as sensitive group, partial response (PR) and no response (NR) as refractory or relapsed (RR) group, b) the in vitro treatment with daunorubicin or not into 4 groups, i.e. CR, CR_D, RR, RR_D. All input peak summits from MACS2 calling were identically extended by 250 bp on either side to a final width of 501 bp and the non-overlapped, merged peak set was identified using the iterative overlap approach(36) (<https://github.com/corceslab/ATAC_IterativeOverlapPeakMerging>) which results in the union set of 60,881 reproducible peaks.

**Reads-in-peaks normalization of AML ATAC-seq signal tracks**

The ATAC-seq signal tracks were generated by the reads-in-peaks normalization approach(37) as the patient’s primary sample of varying quality was not comparable for the varying signal-to-noise ratio, i.e. fraction of mapped reads in called peaks (FRiP). Briefly, the genome was segmented into 100 bp bins and the Tn5 offset-corrected ATAC-seq reads falling in bins overlapping peak regions and background regions were quantified separately. The ratio between peak reads sum and background reads sum was calculated as the scale factor to normalize each sample read depth in peak regions or background into the constant total 30 million reads per sample. The bins of read counts for each sample were converted into bigwig files using bedGraphToBigWig(14).

**Construction of ATAC-seq peak accessibility matrix**

To quantify the Tn5 insertions within the merged peak set for each sample, first the BAM files for 30 samples were identically converted to BEDPE format and adjusted for the Tn5 offset (“+” stranded +4 bp, “-” stranded -5 bp). Then the number of Tn5 cleavage falling in each peak was counted for each sample which was collapsed into a 60,881×30 peaks-by-samples count matrix. The raw count matrix was normalized first using log_2_(CPM) with the prior count of 5 and then quantile normalized using the normalize.quantiles() function in R package preprocessCore. The PCA analysis was performed using the prcomp() function in R package ggfortify.

**Distal binarization identifies daunorubicin-responsive peaks**

To identify differential peaks that are unique to multiple groups, we applied the classification method termed “distal binarization” as previously described(37). Briefly, the intra-group mean 𝜇*_i,j_* and intra-group standard deviation 𝜎*_i,j_* of normalized ATAC-seq signal in each peak *i* across all the samples in each group *j* were calculated. For each peak *i*, the groups were ranked by the intra-group mean (𝜇*_i,j_^1st^*, 𝜇*_i,j_^2nd^*,..., 𝜇*_i,j_^kth^*,...,𝜇*_i,j_^Nth^*) and iterated from the group *j^1st^* with *1st* lowest mean ATAC-seq signal. The criteria to define the break point is if the next-lowest intra-group mean 𝜇*_i,j_^(k+1)th^* of the next group *(k+1)th* is greater than the maximum of other *k* lower intra-group means plus their intra-group standard deviations *max*(𝜇*_i,j_^1st^* + 𝜎*_i,j_^1st^* , …, 𝜇*_i,j_^kth^* + 𝜎*_i,j_^kth^*). A sign matrix was assigned with “0” before break point and “1” after break point. To assess the statistical significance of the binary assigned group-unique peaks, the limma’s eBayes differential test was performed on the normalized ATAC-seq signal matrix with the contrast matrix input which is defined as the binary assigned differential groups combination. The binary assigned group-unique peaks below the FDR cutoff of 0.05 were kept and sorted based on the binary assigned differential groups combinations. The distal binarization method results in 14,260 significant group-unique peaks. The sorted ATAC-seq signal matrix of binary assigned group-unique peaks was plotted using the R package ComplexHeatmap(38). Daunorubicin-responsive ATAC-seq peaks were identified based on the binary assigned group combinations, for examples, peaks which were assigned to the combination of “CR:1, CR_D:1, RR:1, RR_D:0” were extracted as the refractory/relapsed AML patients-unique daunorubicin-deactivated ATAC-seq peaks.

**Motif enrichment analysis in daunorubicin-responsive peaks**

The motifs enrichment analysis within group-unique peak sets was performed first using the R package motifmatchr to compute the occurrence of known motifs from CIS-BP database. To identify the motif specifically enriched within the group-unique peak sets, the background motif occurrences within the total merged ATAC-seq peak set were computed as well. The hypergeometric test of the motif representation within the group-unique peak sets compared to the total merged peak set was calculated using the R package phyper with the cutoff of P-value < 10^-5^. The motif enriched in at least one but not all groups were sorted based on log_10_(P-value) and plotted using the R package ComplexHeatmap(38).

**Correlation analysis to predict daunorubicin-responsive peak-to-gene links**

To infer the causal regulatory links between daunorubicin-responsive ATAC-seq peaks and target genes, we performed the correlation analysis between chromatin accessibility and gene expression using 20 cases of paired RNA-seq and ATAC-seq data from primary AML patients as previously described(37). First, the ATAC-seq peak accessibility count matrix was normalized using log_2_(CPM) with the prior count of 5 and the RNA-seq gene expression count matrix was normalized using log_2_(RPKM+1). Second, the less variable (below 25th quantile) ATAC-seq peaks and genes were filtered from both matrices as they will arise false-positive predictions based on the correlation approach. Third, all the potential interactions between ATAC-seq peaks and gene promoters (within -1500 to +500 bp from the TSSs) within 0.5 Mbp were extracted and the Pearson correlation was calculated for each potential interaction. To assess the statistical significance of identified correlations between ATAC-seq peak accessibility and gene expression, the null distribution was constructed on each single chromosome by correlating the chromatin accessibility of random 10000 peaks from the other different chromosomes and the gene expression of each gene in this chromosome. The correlation model of non-specific interactions between ATAC-seq peak chromatin accessibility and gene expression on a single chromosome were estimated by the mean and standard deviation calculated from the “trans” peak-to-gene interactions. The cutoff of p-value below 0.05 was applied to filter the non-significant correlations. The peak-to-gene links falling within the gene promoter regions were filtered to identify potential long-range interactions between distal regulatory elements (e.g. CTCF loops) and target genes. This results in 32,060 significant peak-to-gene links in total. To identify the daunorubicin-responsive peak-to-gene links, the group-unique differential peaks from distal binarization were overlapped with the predicted peak-to-gene links. The potential interactions driven by CTCF from daunorubicin-responsive peak-to-gene links were further extracted by overlapping the CTCF motif scanned by FIMO with the cutoff of p-value below 10^-5^. The predicted peak-to-gene links were visualized using the R package plotgardener(20).
