## Supplementary Figures for "Targeting the 3D genome by anthracyclines for chemotherapeutic effects"

[7]Lead Contact

### Supplementary Figure 1

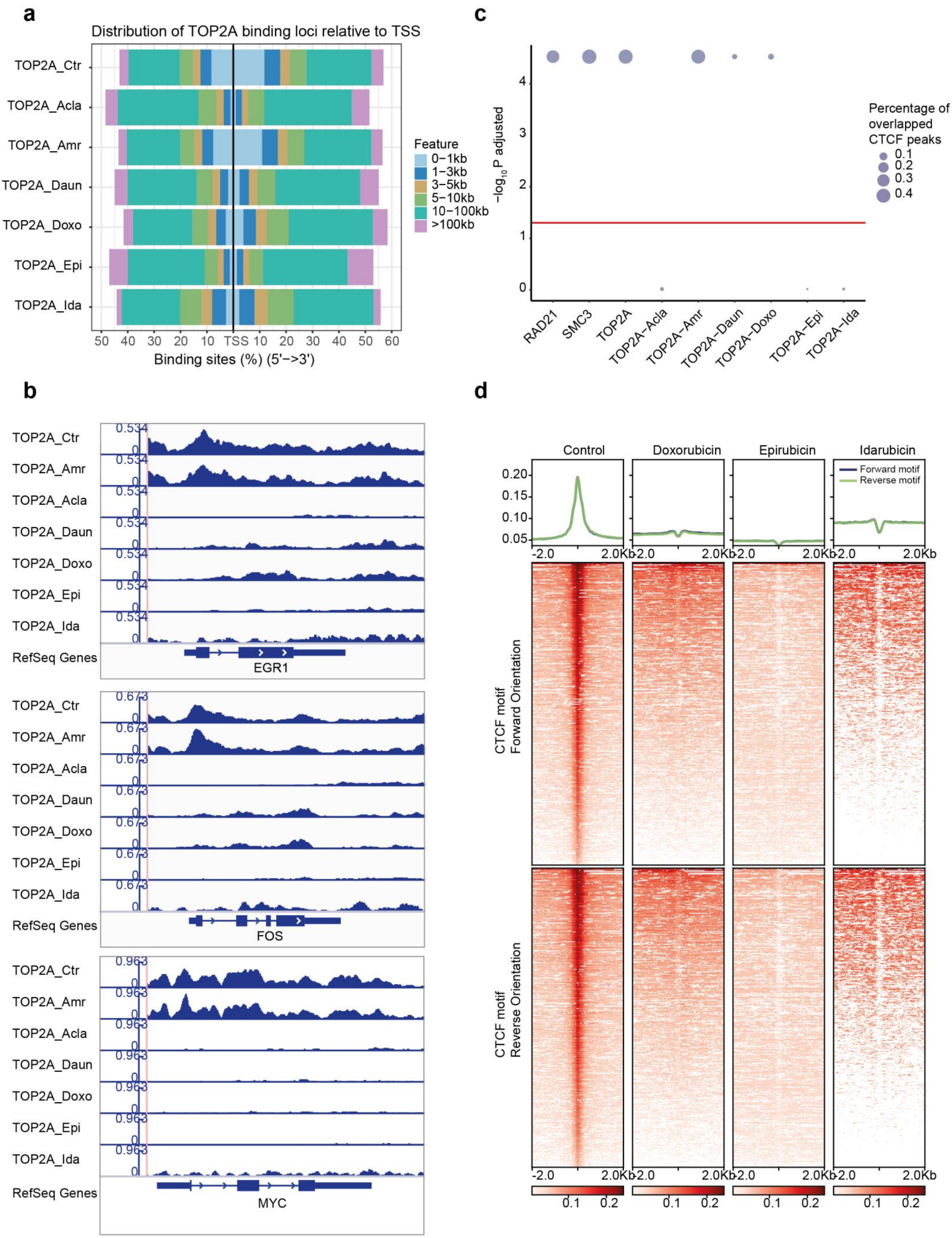

**Figure S1. Genomic patterns of redistributed TOP2A by anthracyclines.**

**(a)** The distribution patterns of TOP2A relative to transcriptional start site (TSS) after different anthracyclines treatments. From top to bottom, untreated K562 cells (Ctr), cells treated with Acla, Amr, Daun, Doxo, Epi and Ida (10 µM for 2 hours) are shown.

**(b)** Snapshot of TOP2A ChIP-seq from untreated K562 cells (Ctr) and K562 cells treated with Amr, Acla, Daun, Doxo, Epi and Ida at immediate early genes (IEGs) loci. From top to bottom, IEG loci of *EGR1*, *Fos* and *Myc* are shown.

**(c)** Association of TOP2A peaks from untreated K562 and TOP2A peaks from Acla-, Amr-, Daun-, Doxo-, Epi- and Ida-treated K562 cells with published CTCF ChIP-seq peaks. The cohesin subunits RAD21 and SMC3 were tested as positive control of CTCF associated factors. The *y* axis indicates the significance of the enrichment of TOP2A within CTCF peaks. The *P*-values were calculated by permutation test ($n$=50,000) and multiple testing correction by the Benjamini-Hochberg approach. The red horizontal line indicates the cutoff of *P*-value 0.05. Circle size indicates the ratio of CTCF peaks covered by the respective TOP2A peaks (scale, 0-1).

**(d)** TOP2A ChIP-seq profile around chromatin looping anchors from untreated and Doxo-, Epi- and Ida-treated samples. Chromatin looping anchors were defined by co-binding sites of CTCF and cohesin subunit RAD21 from ChIP-seq experiments. CTCF motifs in chromatin looping anchors were identified by FIMO with the threshold of *P*-value $< 10^{-5}$ and categorized as forward or reverse orientation based on their detected DNA strand. Normalized TOP2A ChIP-seq reads signal were shown within ± 2 kbp from CTCF motif center.

### Supplementary Figure 2

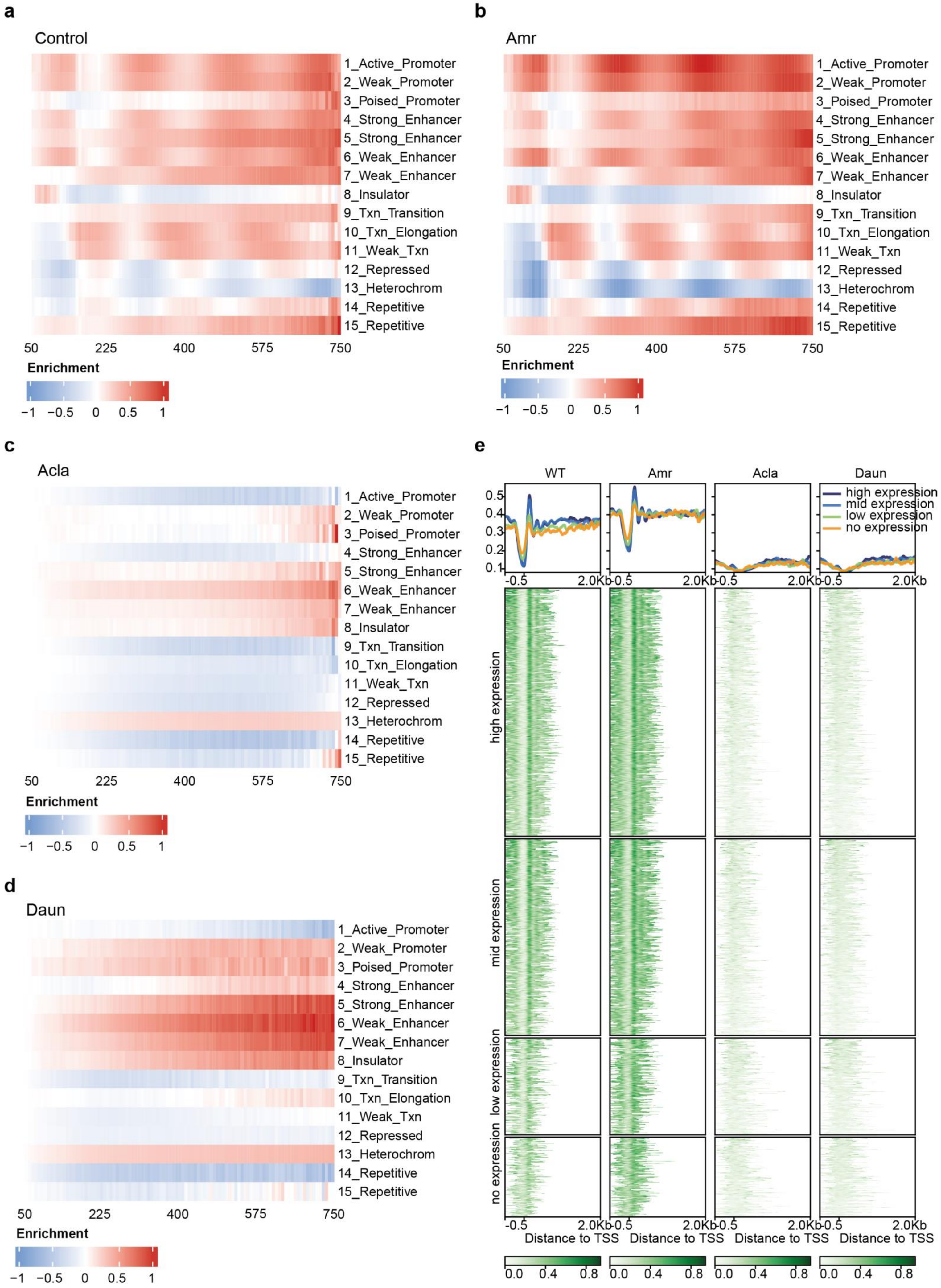

**Figure S2. The chromatin structure changes at different chromatin states after anthracycline treatments.**

**(a-d)** Normalized ATAC-seq fragments enrichment from untreated (**a**) and Amr- (**b**), Acla- (**c**) and Daun-treated samples (**d**) in 15 classes of chromatin states predicted by chromHMM in K562 cells. All drug-treated ATAC-seq samples were uniformly treated with 10 µM of the compound for 2 hours. The Tn5 insertion size ranging from 50 bp to 750 bp were divided into 100 bins and the percent of insertions in each bin were normalized to the maximal percent within each state. The enrichment of Tn5 insertions in different-size bins were calculated by dividing the normalized percent of insertions for each state by the genome-wide normalized percent of insertions. scale: -1 to +1.

**(e)** Nucleosome occupancy around transcription start sites (TSSs) from untreated and Amr-, Acla- and Daun-treated samples. The TSSs of genes were categorized into high-, middle-, low- and no-expression classes based on the RNA-seq data in K562. The nucleosome occupancy profiles called from ATAC-seq by NucleoATAC were shown within - 500 bp to 2 kbp relative to TSSs.

### Supplementary Figure 3

**a**

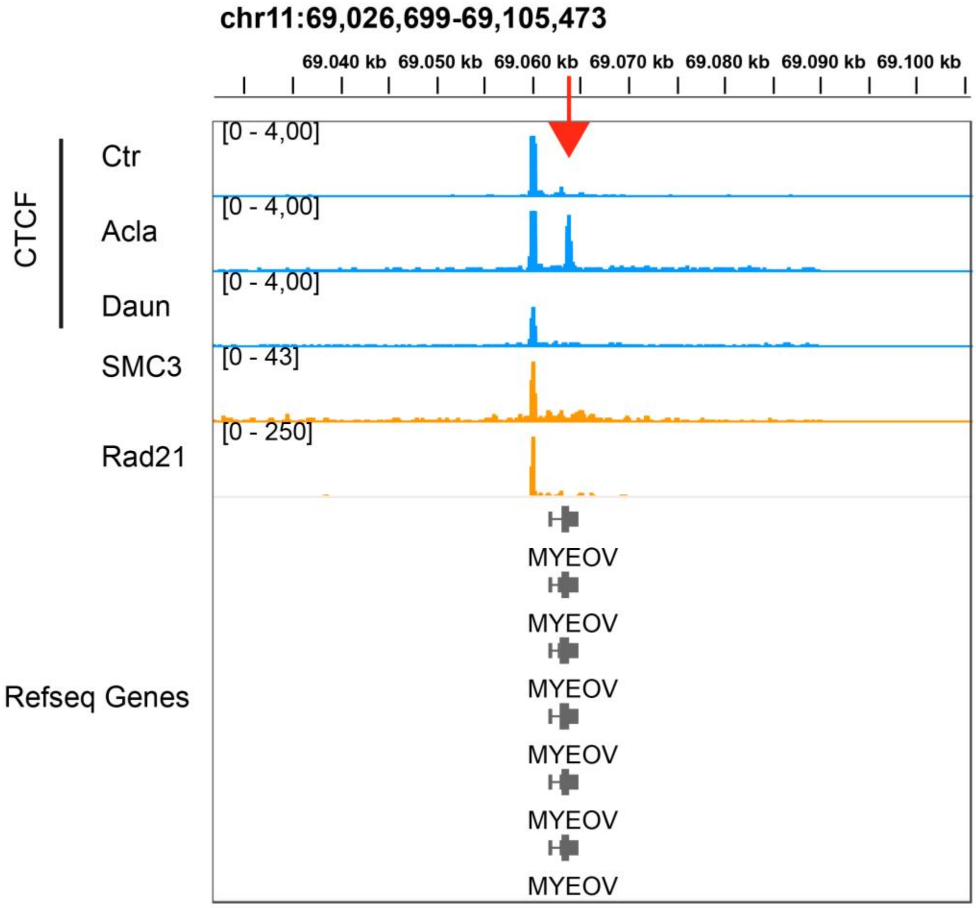


**b**

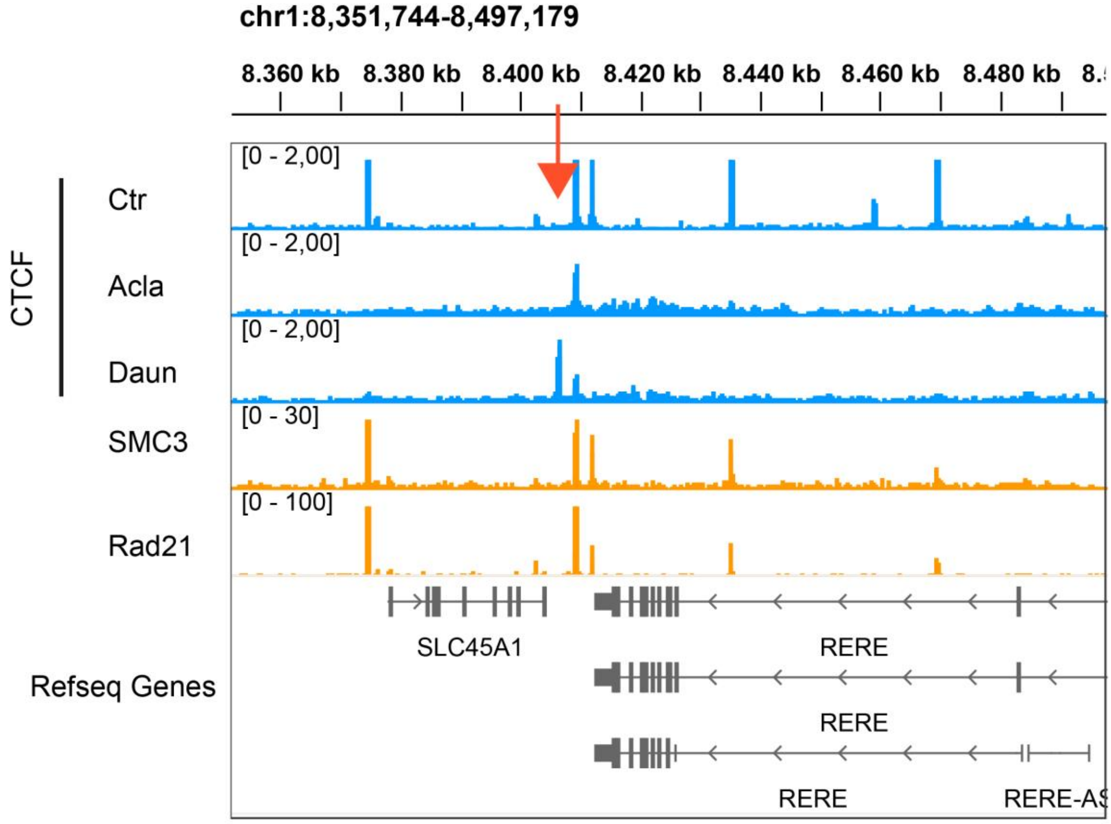


**Figure S3. Anthracyclines redistribute CTCF binding**
**(a-b)** Snapshot of normalized CTCF ChIP-seq signals from untreated, Acla- and Daun-treated K562 cells (10 µM for 2 hours) at RXRA and EPS15L1 loci. ENCODE ChIP-seq data against cohesin subunits RAD21 and SMC3, topologically associating domains (TADs) in K562 cells, and gene annotation are shown. Drug-specific gained CTCF binding sites are indicated by arrows.

#### Supplementary Figure 4

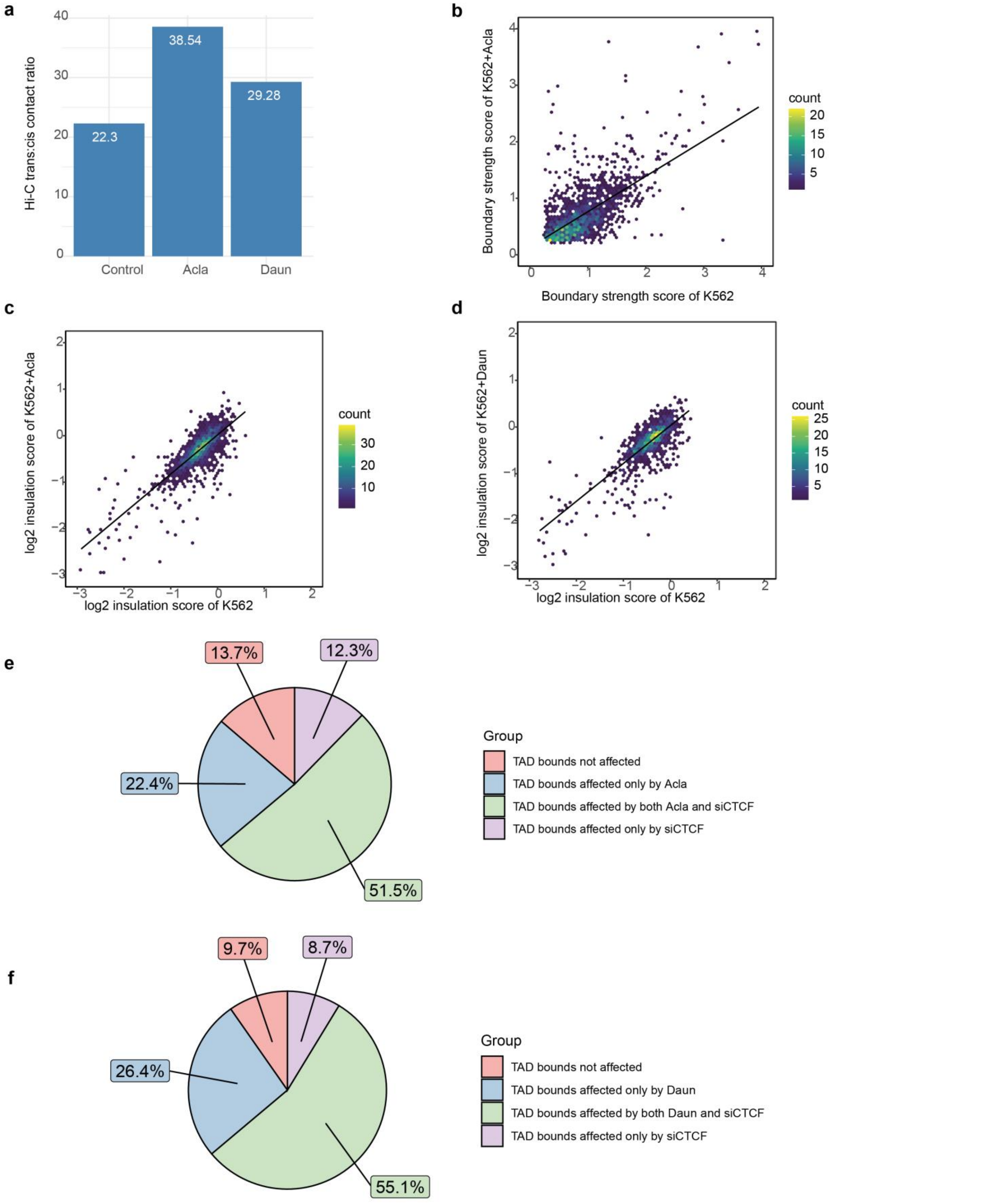


**Figure S4. Anthracyclines disrupt TAD boundaries and promote trans contacts**

**(a)** Barplot of ratios of inter- vs. intra-chromosomal Hi-C contact reads in Control, Acla- and Daun-treated K562 cells (10 µM for 2 hours).

**(b)** Scatterplot of TAD boundary strength scores identified in Control (*x* axis) Acla-treated (*y* axis) Hi-C matrices.

**(c-d)** Scatterplot of log2 insulation scores identified in Control (*x* axis) and Acla- (**c**) and Daun-treated (**d**) (*y* axis) Hi-C matrices.

**(e)** Distribution of TAD boundary changes in response to Acla and siCTCF treatments. Pie chart shows the proportion of TAD boundaries affected under different conditions.

**(f)** Distribution of TAD boundary changes in response to Daun and siCTCF treatments. Pie chart shows the proportion of TAD boundaries affected under different conditions.

### Supplementary Figure 5

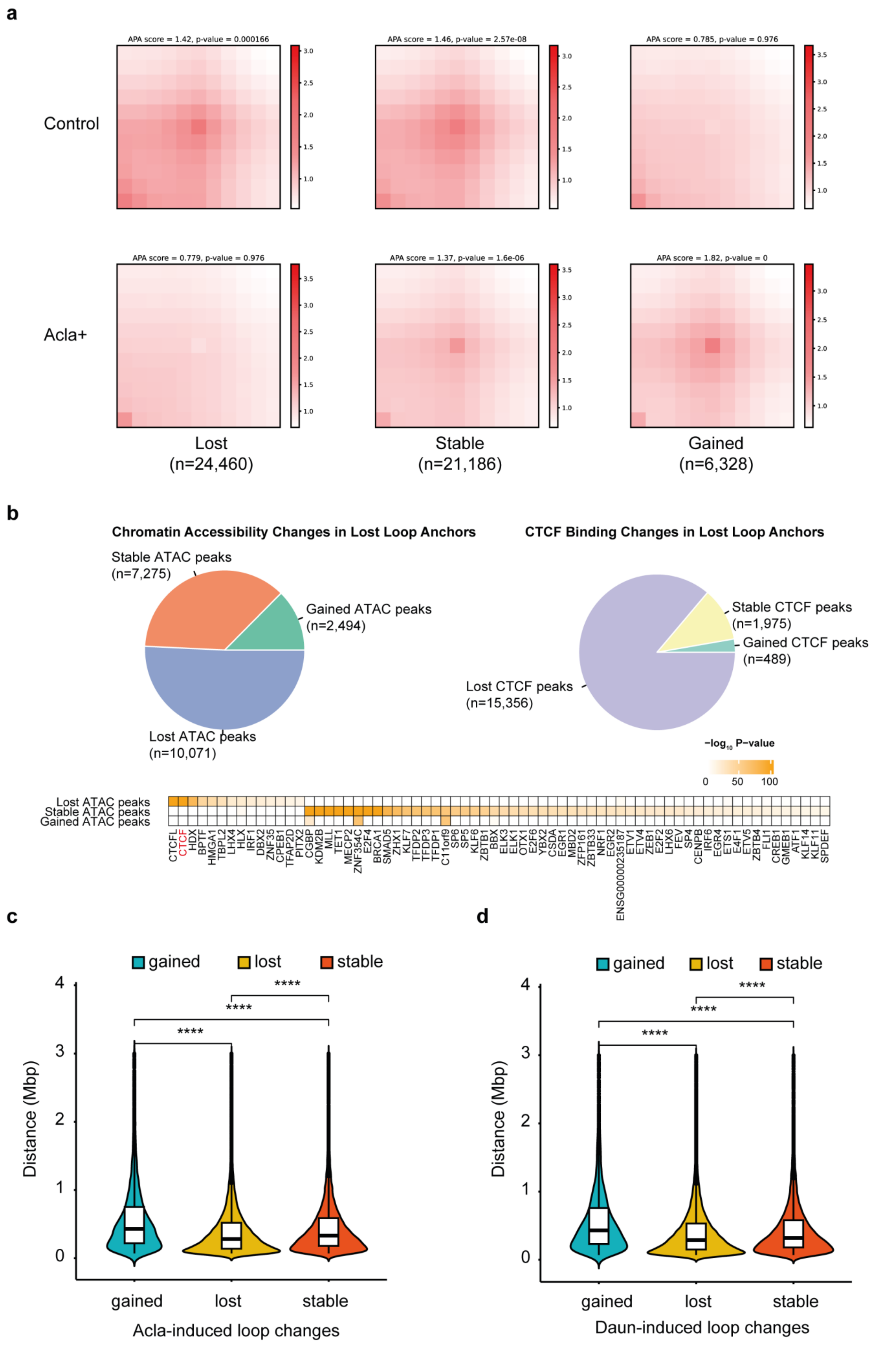

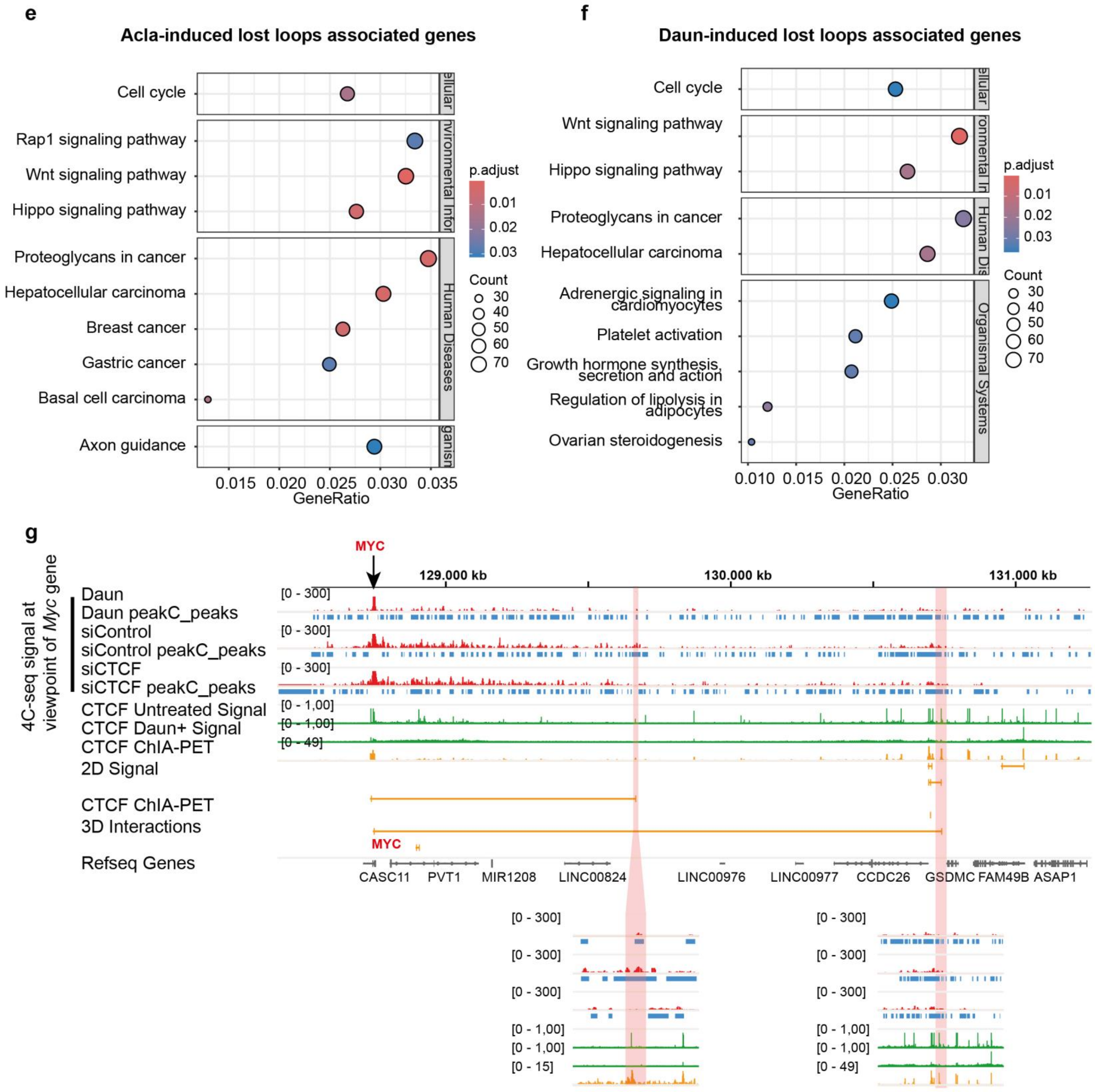
e
Acla-induced lost loops associated genes
Cell cycle
Rap1 signaling pathway
Wnt signaling pathway
Hippo signaling pathway
Proteoglycans in cancer
Hepatocellular carcinoma
Breast cancer
Gastric cancer
Basal cell carcinoma
Axon guidance
ellular
nvironmental Infor
Human Diseases
ganism
0.015 0.020 0.025 0.030 0.035
GeneRatio
p.adjust
0.01
0.02
0.03
Count
30
40
50
60
70
f
Daun-induced lost loops associated genes
Cell cycle
Wnt signaling pathway
Hippo signaling pathway
Proteoglycans in cancer
Hepatocellular carcinoma
Adrenergic signaling in cardiomyocytes
Platelet activation
Growth hormone synthesis, secretion and action
Regulation of lipolysis in adipocytes
Ovarian steroidogenesis
ellular
ronmental In
Human Dis
Organismal Systems
0.010 0.015 0.020 0.025 0.030
GeneRatio
p.adjust
0.01
0.02
0.03
Count
30
40
50
60
70
g
4C-seq signal at viewpoint of Myc gene
MYC
129,000 kb
130,000 kb
131,000 kb
Daun
Daun peakC_peaks
siControl
siControl peakC_peaks
siCTCF
siCTCF peakC_peaks
CTCF Untreated Signal
CTCF Daun+ Signal
CTCF ChIA-PET
2D Signal
CTCF ChIA-PET
3D Interactions
Refseq Genes
[0 - 300]
[0 - 1,00]
[0 - 49]
[0 - 15]
CASC11
PVT1
MIR1208
LINC00824
LINC00976
LINC00977
CCDC26
GSDMC
FAM49B
ASAP1

h

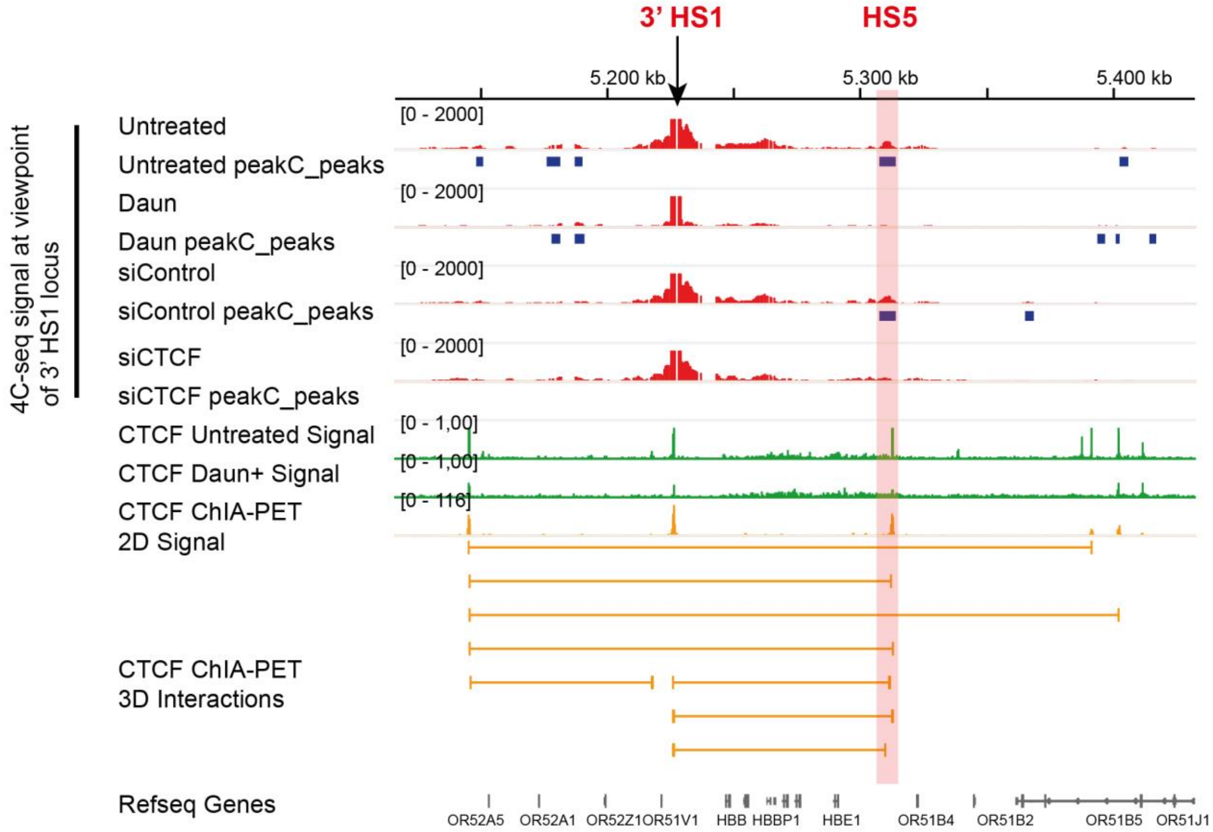

3' HS1
HS5
5,200 kb
5,300 kb
5,400 kb
4C-seq signal at viewpoint of 3' HS1 locus
Untreated
Untreated peakC_peaks
Daun
Daun peakC_peaks
siControl
siControl peakC_peaks
siCTCF
siCTCF peakC_peaks
CTCF Untreated Signal
CTCF Daun+ Signal
CTCF ChIA-PET 2D Signal
CTCF ChIA-PET 3D Interactions
Refseq Genes
[0 - 2000]
[0 - 1,00]
[0 - 116]
OR52A5
OR52A1
OR52Z1
OR51V1
HBB
HBBP1
HBE1
OR51B4
OR51B2
OR51B5
OR51J1


i

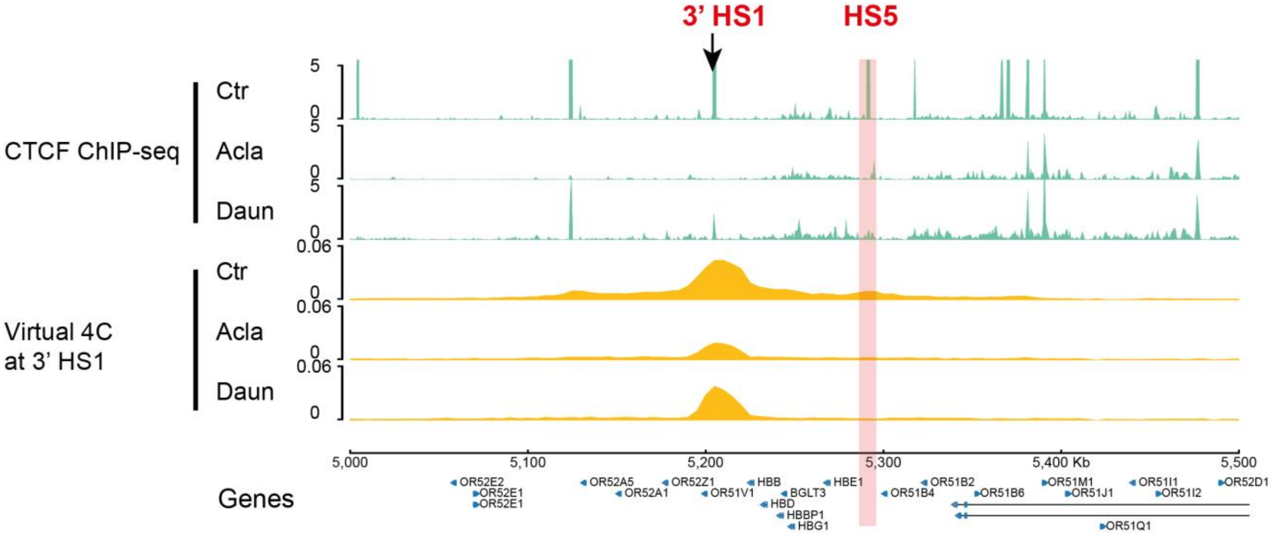

3' HS1
HS5
CTCF ChIP-seq
Ctr
Acla
Daun
Virtual 4C at 3' HS1
0.06
5
0
5,000
5,100
5,200
5,300
5,400 Kb
5,500
Genes
OR52E2
OR52E1
OR52A5
OR52A1
OR52Z1
OR51V1
HBB
HBD
BGLT3
HBBP1
HBG1
HBE1
OR51B4
OR51B2
OR51B6
OR51M1
OR51J1
OR51Q1
OR51I1
OR51I2
OR52D1


j

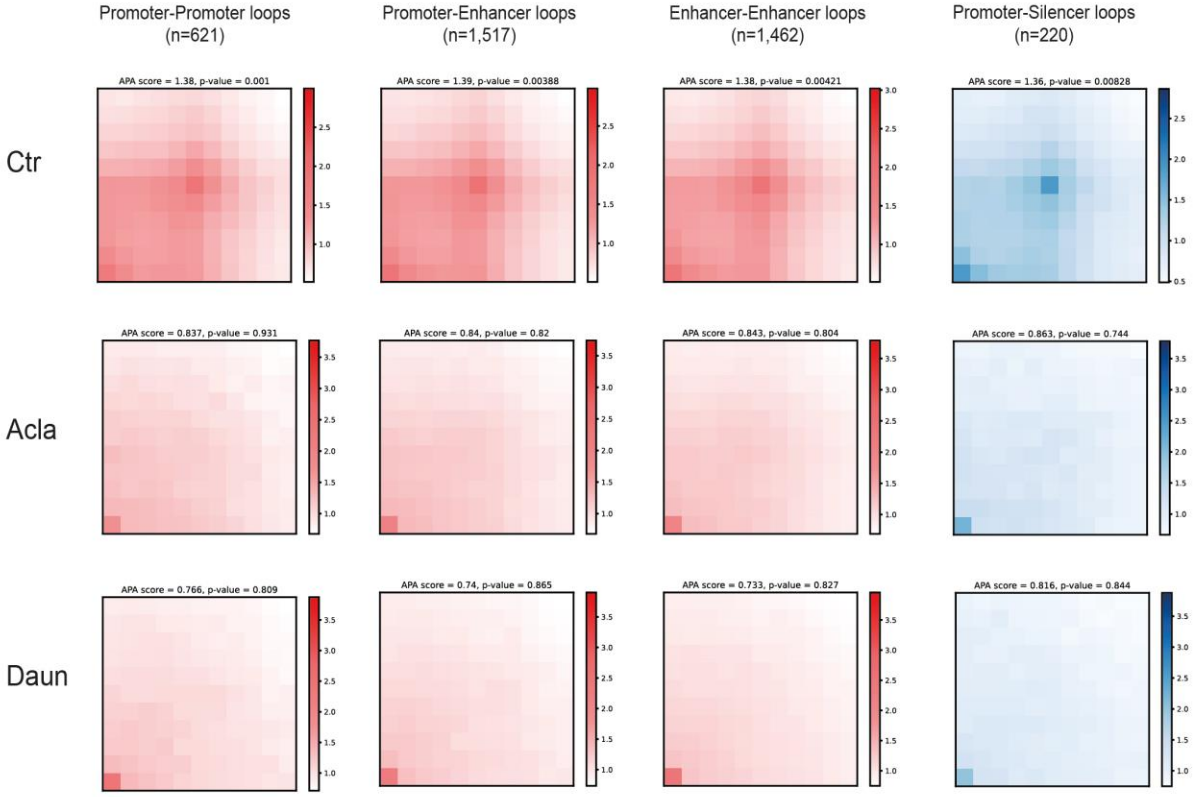

Promoter-Promoter loops (n=621)
Promoter-Enhancer loops (n=1,517)
Enhancer-Enhancer loops (n=1,462)
Promoter-Silencer loops (n=220)
Ctr
Acla
Daun
APA score = 1.38, p-value = 0.001
APA score = 1.39, p-value = 0.00388
APA score = 1.38, p-value = 0.00421
APA score = 1.36, p-value = 0.00828
APA score = 0.837, p-value = 0.931
APA score = 0.84, p-value = 0.82
APA score = 0.843, p-value = 0.804
APA score = 0.863, p-value = 0.744
APA score = 0.766, p-value = 0.809
APA score = 0.74, p-value = 0.865
APA score = 0.733, p-value = 0.827
APA score = 0.816, p-value = 0.844

**Figure S5. Epigenomic signature of disrupted chromatin loops by anthracyclines**
**(a)** Aggregate peak analysis of rewired chromatin loops in Acla-treated K562 cells (gained, lost and stable) within ± 50 kbp window. Differential loops were determined by diffPeakachu when the fold-change of probability score deviates from the Gaussian mixture model with a *P*-value < 0.05.
**(b)** Epigenomic signatures of lost loops in Acla-treated K562 cells. In the upper panel, the pie charts of differential Acla-responsive (gained, lost and stable) ATAC-seq peaks (left) and CTCF ChIP-seq peaks (right) residing in the lost looping anchors were shown. In the lower panel, the motif enrichment in differential Acla-responsive ATAC-seq peaks was shown. *P*-values were calculated using hypergeometric distribution compared with background peaks and significantly enriched TF motifs were determined with the cutoff of *p*-value below 10-10.
**(c-d)** Distribution of distance spanning of different categorial rewired chromatin loops (gained, lost and stable) in Acla- (c) and Daun-treated (d) K562 cells (bp). P-values were calculated using the two-sided Wilcoxon rank-sum test.
**(e-f)** KEGG pathway enrichment of genes associated with Acla- (e) and Daun-induced (f) lost loops. The genes with promoters overlapping anchors of lost loops were identified. The cutoff of adjusted P-values below 0.05 was used.
**(g)** 4C-seq profiles displaying the in cis chromatin contacts of the viewpoint at Myc promoter-proximal CTCF site. From top to bottom, tracks of 4C signal and peaks called by peakC in Daun-, si-Ctr- and si-CTCF-treated 10 milions K562 cells, CTCF ChIP-seq signal of untreated and Daun-treated K562 cells, CTCF ChIA-PET 2D signal and 3D interactions are shown. The viewpoint region of Myc gene is indicated by arrow. The zoomed-in views of the two interacted partner CTCF sites are provided as highlighted by red bar.
**(h)** 4C-seq profiles displaying the in cis chromatin contacts of the viewpoint at 3’ HS1 CTCF site in β-globin locus. From top to bottom, tracks of 4C signal and peaks called by peakC in untreated, Daun-, si-Ctr- and si-CTCF-treated 10 milions K562 cells, CTCF ChIP-seq signal of untreated and Daun-treated K562 cells, CTCF ChIA-PET 2D signal and 3D interactions are shown. The viewpoint region of 3’ HS1 site is indicated by arrow. The interacted partner HS5 CTCF site are highlighted by red bar.
**(i)** Virtual 4C analysis from the Hi-C data with the viewpoint at the 3’ HS1 CTCF element at β-globin locus. The viewpoint anchored at 3’ HS1 CTCF-bound site is indicated by arrow. The interacted HS5 CTCF-bound partner site is indicated by red bar. Normalized CTCF ChIP-seq and virtual 4C signal tracks from untreated (Ctr), Acla- and Daun-treated K562 cells are shown.
**(k)** Aggregate peak analysis of promoter-promoter, enhancer-promoter, enhancer-enhancer and putative silencer-promoter loops within ± 50 Kbp window. From top to bottom: Control, Acla- and Daun-treated K562 cells were shown. The promoter-promoter loops were defined as both anchors overlapping with H4K3me3-marked promoters within (-1500bp, +500bp) relative to TSS. The enhancer-enhancer loops were defined as both anchors overlapping H3K27ac-marked enhancers which were excluded from promoter annotation. The enhancer-promoter loops were defined as two anchors overlapping with H4K3me3-marked promoter and H3K27ac-marked enhancer respectively. The silencer-promoter loops were defined as two anchors overlapping with silencers from high-throughput screening and promoters within (-1500bp, +500bp) relative to TSS respectively.

### Supplementary Figure 6

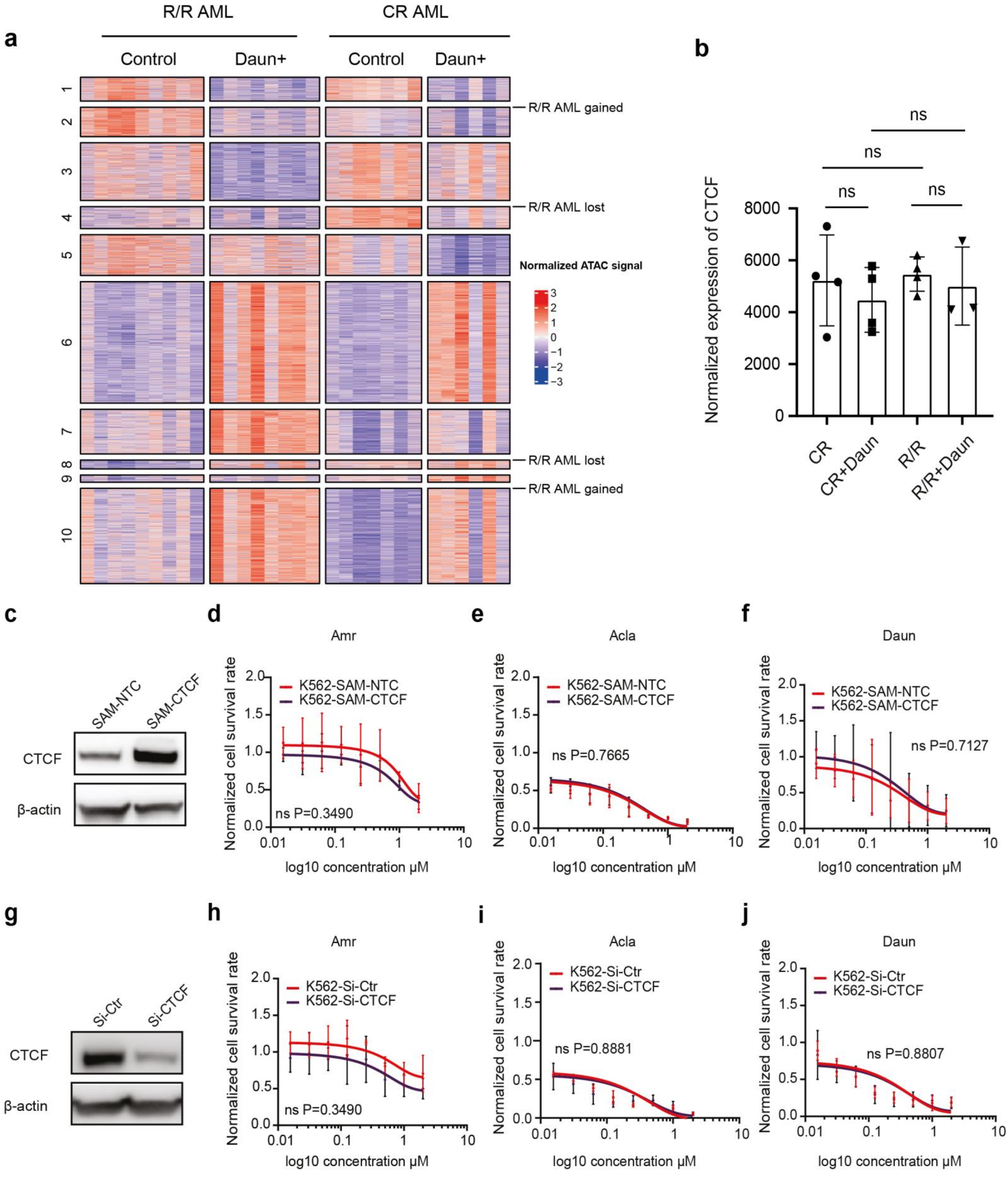

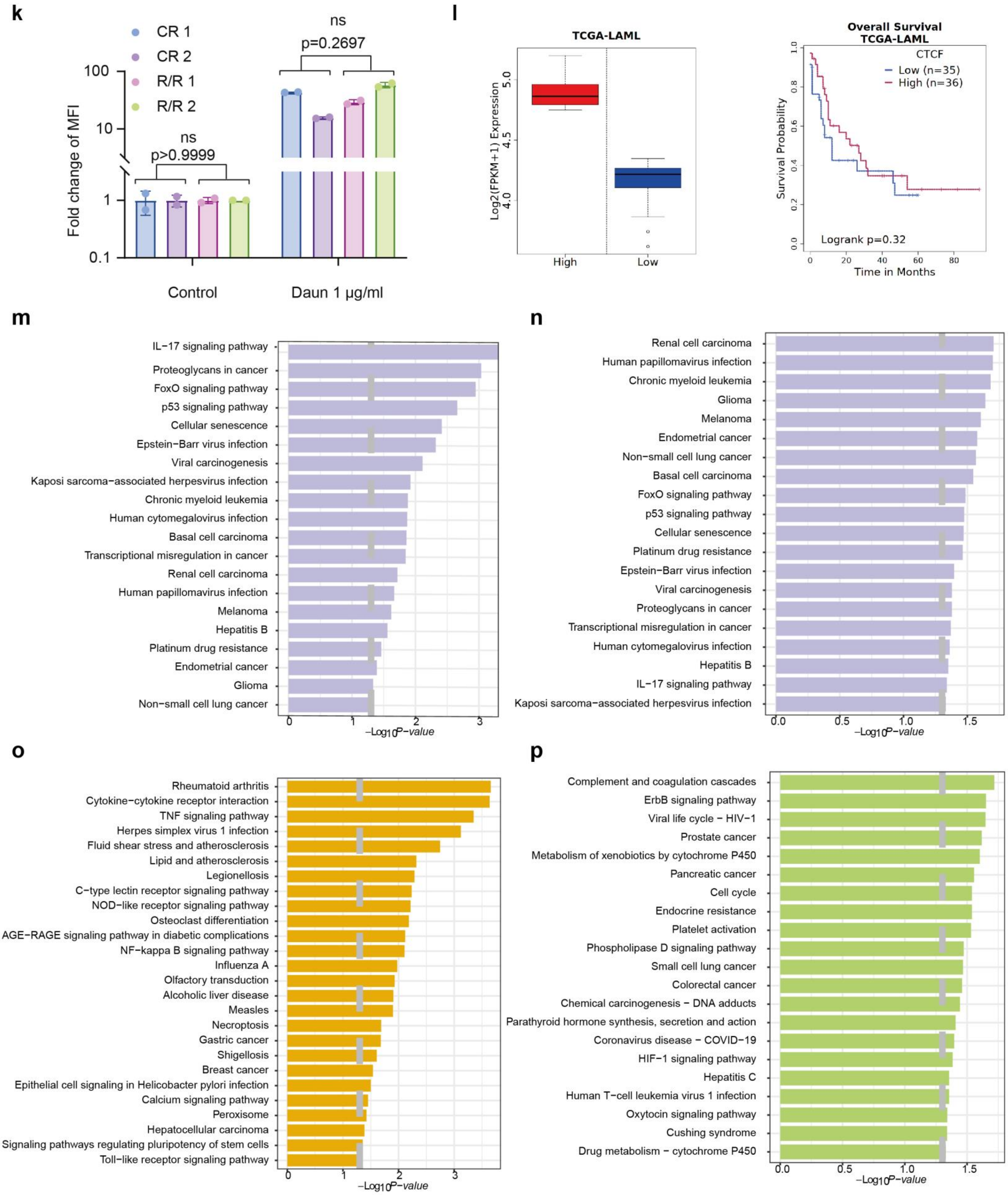

k
CR 1
CR 2
R/R 1
R/R 2
ns
p=0.2697
ns
p>0.9999
Fold change of MFI
100
10
1
0.1
Control
Daun 1 μg/ml
l
TCGA-LAML
Log2(FPKM+1) Expression
5.0
4.5
4.0
High
Low
Overall Survival
TCGA-LAML
CTCF
Low (n=35)
High (n=36)
Survival Probability
Logrank p=0.32
Time in Months
0
20
40
60
80
m
IL−17 signaling pathway
Proteoglycans in cancer
FoxO signaling pathway
p53 signaling pathway
Cellular senescence
Epstein−Barr virus infection
Viral carcinogenesis
Kaposi sarcoma−associated herpesvirus infection
Chronic myeloid leukemia
Human cytomegalovirus infection
Basal cell carcinoma
Transcriptional misregulation in cancer
Renal cell carcinoma
Human papillomavirus infection
Melanoma
Hepatitis B
Platinum drug resistance
Endometrial cancer
Glioma
Non−small cell lung cancer
0
1
2
3
−Log10P−value
n
Renal cell carcinoma
Human papillomavirus infection
Chronic myeloid leukemia
Glioma
Melanoma
Endometrial cancer
Non−small cell lung cancer
Basal cell carcinoma
FoxO signaling pathway
p53 signaling pathway
Cellular senescence
Platinum drug resistance
Epstein−Barr virus infection
Viral carcinogenesis
Proteoglycans in cancer
Transcriptional misregulation in cancer
Human cytomegalovirus infection
Hepatitis B
IL−17 signaling pathway
Kaposi sarcoma−associated herpesvirus infection
0.0
0.5
1.0
1.5
−Log10P−value
o
Rheumatoid arthritis
Cytokine−cytokine receptor interaction
TNF signaling pathway
Herpes simplex virus 1 infection
Fluid shear stress and atherosclerosis
Lipid and atherosclerosis
Legionellosis
C−type lectin receptor signaling pathway
NOD−like receptor signaling pathway
Osteoclast differentiation
AGE−RAGE signaling pathway in diabetic complications
NF−kappa B signaling pathway
Influenza A
Olfactory transduction
Alcoholic liver disease
Measles
Necroptosis
Gastric cancer
Shigellosis
Breast cancer
Epithelial cell signaling in Helicobacter pylori infection
Calcium signaling pathway
Peroxisome
Hepatocellular carcinoma
Signaling pathways regulating pluripotency of stem cells
Toll−like receptor signaling pathway
0
1
2
3
−Log10P−value
p
Complement and coagulation cascades
ErbB signaling pathway
Viral life cycle − HIV−1
Prostate cancer
Metabolism of xenobiotics by cytochrome P450
Pancreatic cancer
Cell cycle
Endocrine resistance
Platelet activation
Phospholipase D signaling pathway
Small cell lung cancer
Colorectal cancer
Chemical carcinogenesis − DNA adducts
Parathyroid hormone synthesis, secretion and action
Coronavirus disease − COVID−19
HIF−1 signaling pathway
Hepatitis C
Human T−cell leukemia virus 1 infection
Oxytocin signaling pathway
Cushing syndrome
Drug metabolism − cytochrome P450
0.0
0.5
1.0
1.5
−Log10P−value

**Figure S6. Chromatin accessibility-based AML subtyping and dosage of CTCF does not correlate with responses to Daun.**
**(a)** Heatmap of chromatin accessibility signal in AML patient-specific Daun-responsive ATAC peak regions (activated or suppressed). ATAC-seq analyses were performed on AML blasts treated with 1 µg/ml daunorubicin (Daun) for 3 hours. CR: complete remission AML patients; R/R: refractory/relapsed AML patients. The distal binarization method was performed to identify patient-specific Daun-responsive ATAC peaks. The limma eBayes test was performed to determine the statistical significance of differential ATAC peaks between sample groups. The threshold of FDR below 0.05 was used to capture the group-specific significant peaks.
**(b)** Normalized expression level sof CTCF from RNA-seq of the AML patients included in this study before and after Daun treatment (1 µg/ml, 3 hours). Bars show mean value ± s.e.m. $^{ns}p>0.05$., calculated using student's t-test.
(**c**) Overexpression of CTCF in K562 cells using the CRISPRa-SAM system. K562 cells were transduced with the lenti-virus expressing guide RNA targeting the CTCF promoter. A non-targeting guide RNA (SAM-NTC) was used as a negative control. CTCF protein levels were assessed by Western blotting, and β-actin was used as a loading control.
(**d-f**) CellTiter-Blue assay was used to quantify the CTCF-overexpression cell viability. Cells were exposed to a serial dilution of Amr, Acla, and Daun for 24 h, then the live cells were measured. Bars show mean value ± s.e.m. (n =3). ns (versus the Ctr), calculated using 2-way ANOVA.
**(g)** Downregulation of CTCF in K562 cells using RNA interference (RNAi). K562 cells were transfected with small interfering RNAs (si-CTCF) targeting CTCF. A non-targeting siRNA was used as a negative control (si-Ctr). CTCF protein levels after 72h transfection were assessed by Western blotting, with β-actin used as a loading control (n=3).
**(h-j)** CellTiter-Blue assay was used to quantify the CTCF-knockdown cell viability. Cells were exposed to a serial dilution of Amr, Acla, and Daun for 24 h, then the live cells were measured. Bars show mean value ± s.e.m. (n =3). $^{ns}p>0.05$(versus the Ctr), calculated using 2-way ANOVA.
**(k)** FACS is used to quantify the uptake of Daun in AML blast cells. Cells were treated with Daun at a final concentration of 1 µg/ml for 3 h. Then fluorescence intensity of the Daun was quantified by FACS. $^{ns}p > 0.05$ (CR versus R/R), calculated using multiple unpaired t tests.
**(l)** Kaplan–Meier analysis of overall survival in TCGA-LAML AML patients stratified by CTCF expression levels. Left: boxplot shows normalized CTCF expression in high- and low-expression groups. Right: red line indicates survival of high CTCF expression group (n = 36); blue line, low expression group (n = 35). P-value calculated using the log-rank test.
**(m-p)** KEGG pathway enrichment analysis based on RNA-seq of R/R AML and CR AML samples after Daun treatment. The common enriched KEGG pathways by Daun treatment were shown for R/R AML in (**m**) and CR AML in (**n**) The R/R AML-specific and CR AML-specific enriched KEGG pathways were shown in (**o**) and (**p**) respectively.

### Supplementary Figure 7

**a**

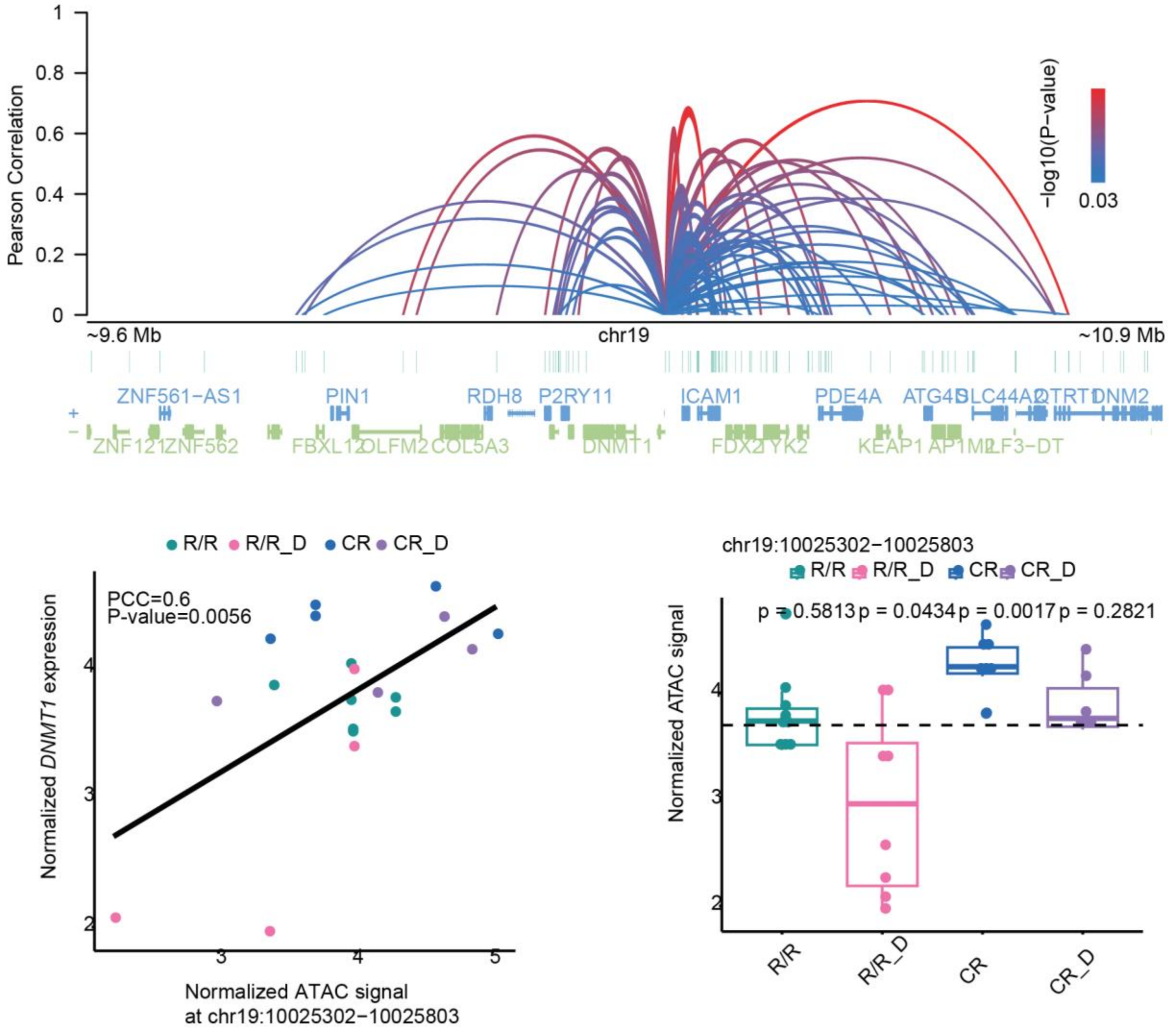


**b**

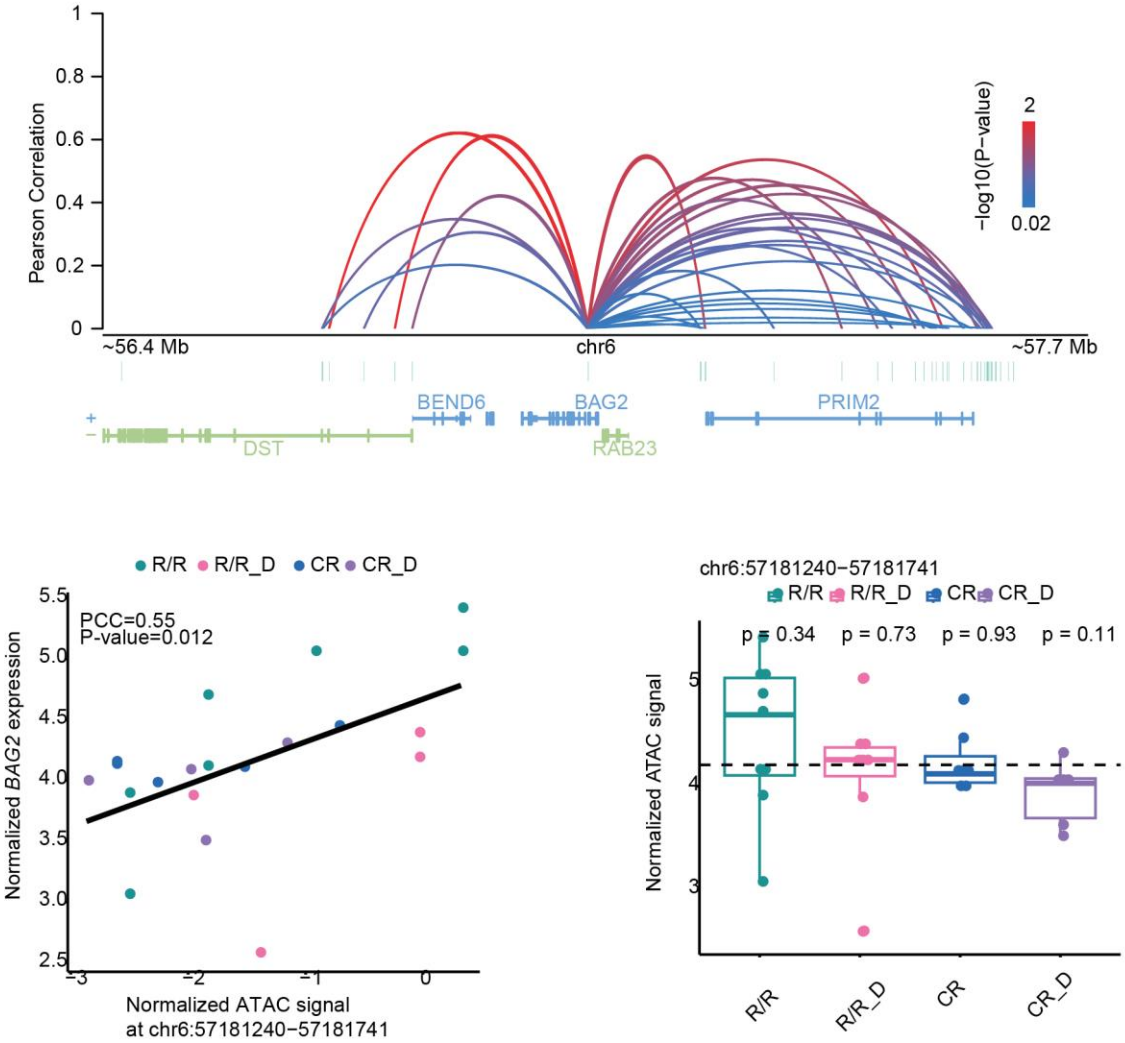

c

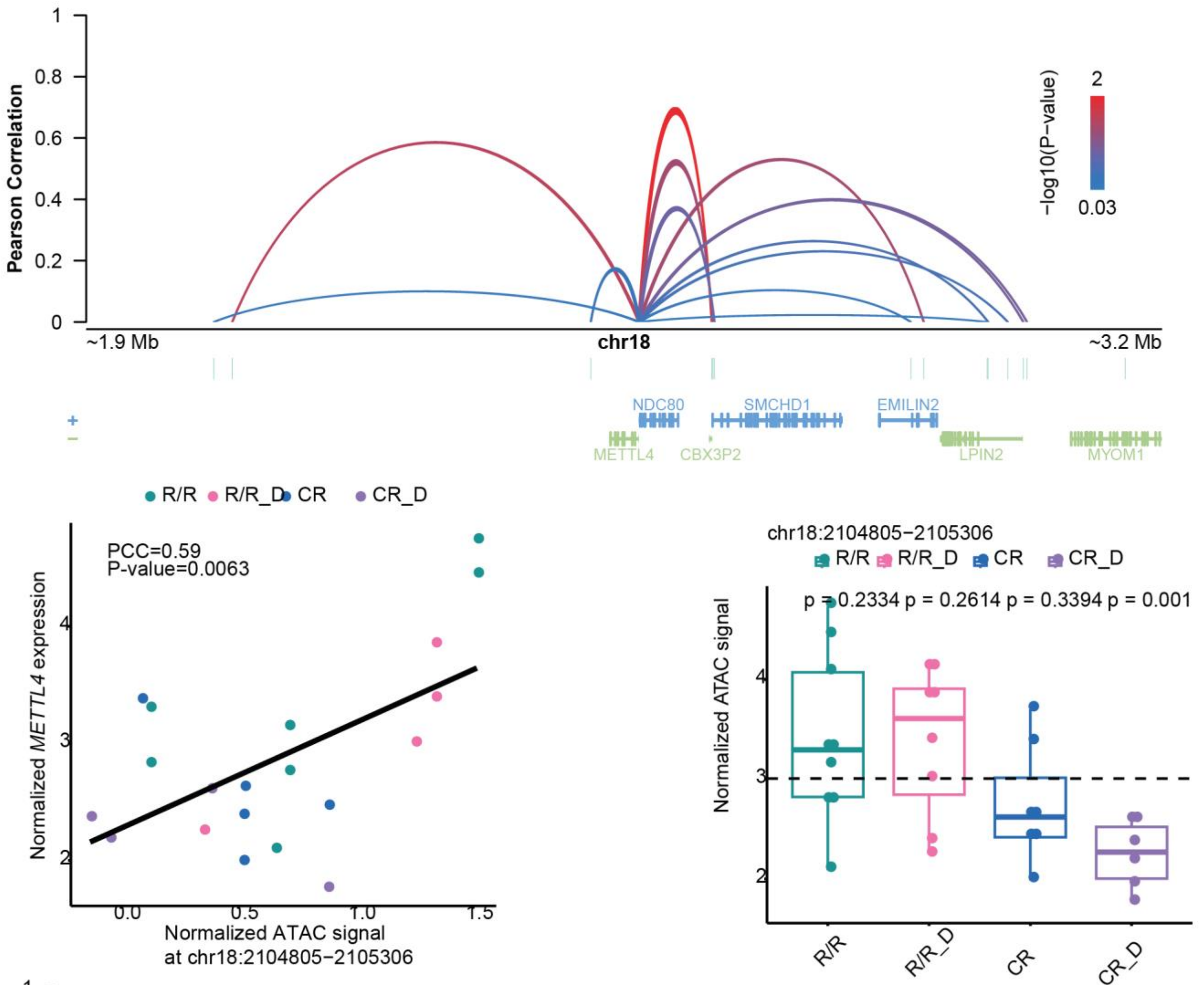

Pearson Correlation
−log10(P−value)
2
0.03
~1.9 Mb
chr18
~3.2 Mb
NDC80
SMCHD1
EMILIN2
METTL4
CBX3P2
LPIN2
MYOM1
R/R R/R_D CR CR_D
PCC=0.59
P-value=0.0063
Normalized METTL4 expression
Normalized ATAC signal
at chr18:2104805−2105306
chr18:2104805−2105306
R/R R/R_D CR CR_D
p = 0.2334 p = 0.2614 p = 0.3394 p = 0.001
Normalized ATAC signal
R/R
R/R_D
CR
CR_D


d

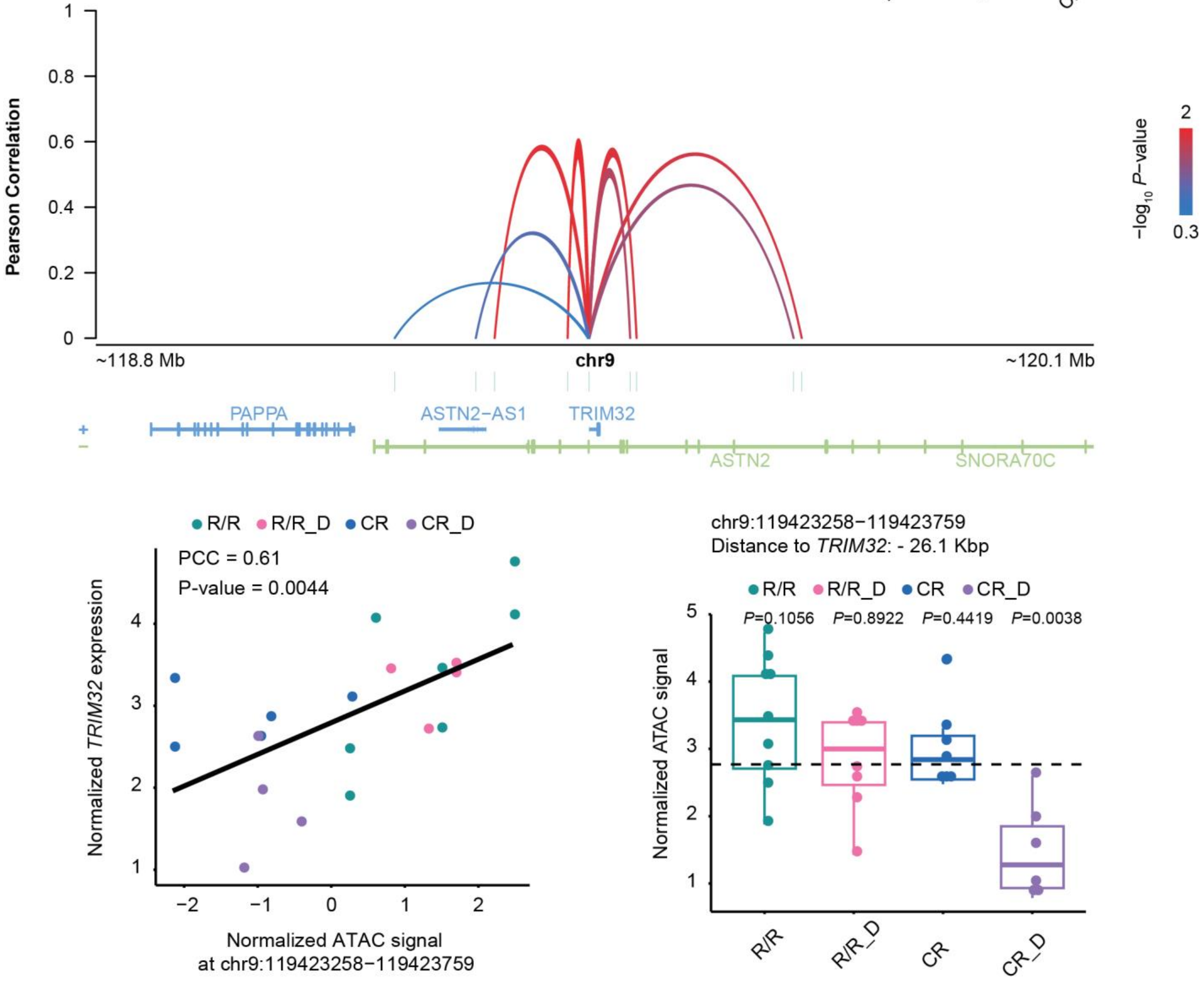

Pearson Correlation
−log10 P−value
2
0.3
~118.8 Mb
chr9
~120.1 Mb
PAPPA
ASTN2-AS1
TRIM32
ASTN2
SNORA70C
R/R R/R_D CR CR_D
PCC = 0.61
P-value = 0.0044
Normalized TRIM32 expression
Normalized ATAC signal
at chr9:119423258−119423759
chr9:119423258−119423759
Distance to TRIM32: - 26.1 Kbp
R/R R/R_D CR CR_D
P=0.1056 P=0.8922 P=0.4419 P=0.0038
Normalized ATAC signal
R/R
R/R_D
CR
CR_D

e

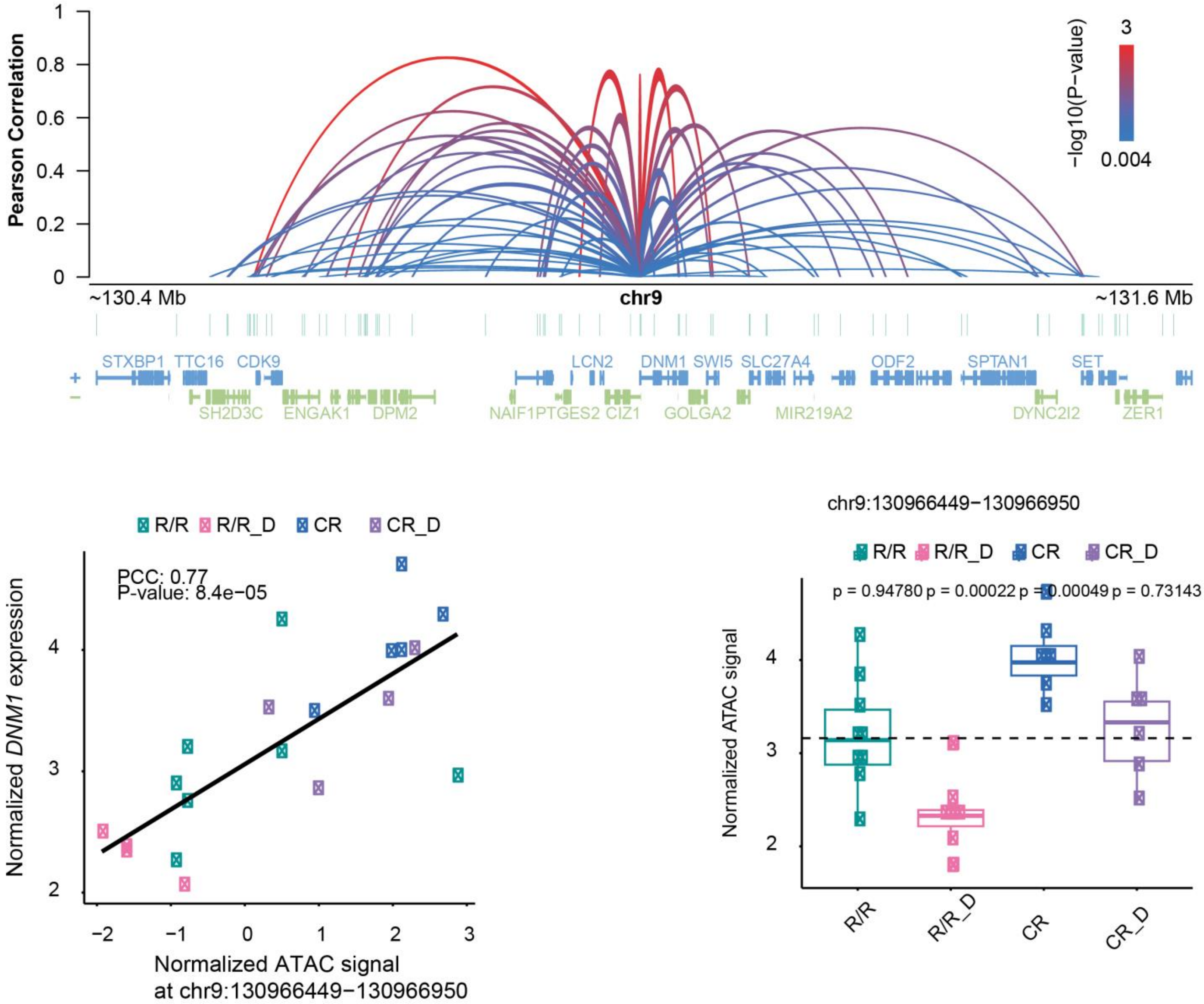

Pearson Correlation
1
0.8
0.6
0.4
0.2
0
−log10(P−value)
3
0.004
~130.4 Mb
chr9
~131.6 Mb
STXBP1 TTC16 CDK9
LCN2 DNM1 SWI5 SLC27A4
ODF2 SPTAN1 SET
SH2D3C ENGAK1 DPM2
NAIF1PTGES2 CIZ1 GOLGA2 MIR219A2
DYNC2I2 ZER1
R/R R/R_D CR CR_D
PCC: 0.77
P-value: 8.4e−05
Normalized DNM1 expression
Normalized ATAC signal
at chr9:130966449−130966950
chr9:130966449−130966950
p = 0.94780 p = 0.00022 p = 0.00049 p = 0.73143
Normalized ATAC signal
R/R
R/R_D
CR
CR_D


f

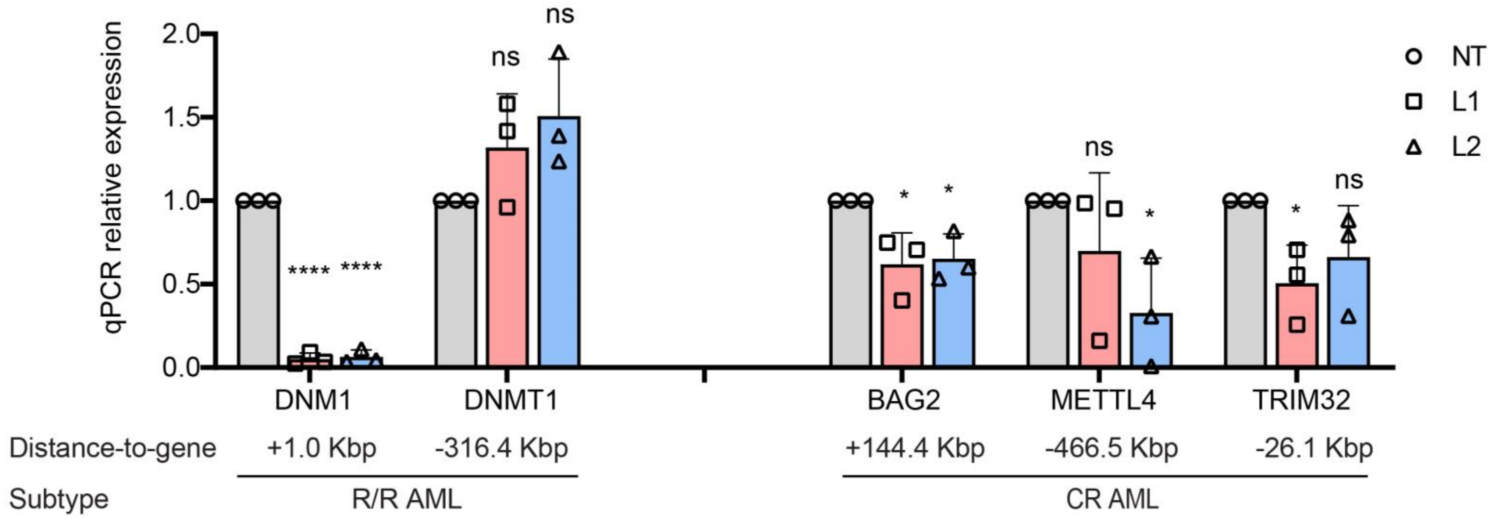

qPCR relative expression
2.0
1.5
1.0
0.5
0.0
ns
****
*
NT
L1
L2
DNM1
DNMT1
BAG2
METTL4
TRIM32
Distance-to-gene
+1.0 Kbp
-316.4 Kbp
+144.4 Kbp
-466.5 Kbp
-26.1 Kbp
Subtype
R/R AML
CR AML

**Figure S7. In silico peak-to-gene linkage analysis to predict susceptible 3D regulatory interactions between CTCF peak and its target gene.**
**(a-e)** Correlation analysis to predict peak-to-gene links where CTCF regulating the target genes: *DNMT1* (**a**), *BAG2* (**b**), *METTL4* (**c**), *TRIM32* (**d**) and *DNM1* (**e**) in the AML subtype-specific fashion. Top: All the potential interactions between target genes and surrounding peaks within the ± 0.5 Mbp window relative to TSS were tested. The Pearson correlation coefficient was calculated to correlate the ATAC-seq accessibility of each peak with the RNA-seq expression of target gene using 20 paired ATAC-seq and RNA-seq samples. The statistical significance was assessed by comparing with the null distribution of correlations between trans-peak-to-gene links. The significant peak-to-gene links were determined using the cutoff of *p*-value below 0.05. Arch height of peak-to-gene links represents the Pearson correlation coefficients. Color indicates the $\log_{10}$ (*p*-value) value. Bottom left: Scatterplot of the ATAC-seq accessibility and RNA-seq gene expression of peak-to-gene link. Color indicates the different patient-treatment groups. Each dot represents an individual sample. Bottom right: Boxplot of the ATAC-seq accessibility of the CTCF motif-containing peak in different sample groups. Color indicates the different patient-treatment groups. Each dot represents an individual sample.
**(f)** Gene expression changes measured by qPCR after CRISPRi targeting CTCF motif-containing peaks predicted to be linked to the susceptible genes to Daun treatment (R/R AML: *DNM1* and *DNMT1*, CR AML: *BAG2*, *TRIM32* and *METTL4*) in K562 cells. NT: dCas9-KRAB with non-targeting genome guide, L1/L2: dCas9-KRAB with 2 different guides targeting the same CTCF motif-containing peak. Error bars represent the standard deviation of 3 biological replicates. ****$P < 0.0001$, *$P<0.05$ and *ns* indicates not significant by two-tailed Student’s t-test.
